# Worldwide cross-ecosystem carbon subsidies and their contribution to ecosystem functioning

**DOI:** 10.1101/271809

**Authors:** Isabelle Gounand, Chelsea J. Little, Eric Harvey, Florian Altermatt

## Abstract

Ecosystems are widely inter-connected by spatial flows of resources^1,2^, yet primarily studied in a local context. Meta-ecosystem models suggest that cross-ecosystem subsidies can play an essential role in ecosystem functioning, notably by controlling local availability of resources for biological communities^3–6^. The general contribution of these resource connections to ecosystem functioning, however, remains unclear in natural systems, due to the heterogeneity and dispersion of data across the ecological literature. Here we provide the first quantitative synthesis on spatial flows of carbon connecting ecosystems worldwide. These cross-ecosystem subsidies range over eight orders of magnitude, between 10^−3^ and 10^5^ gC m^−2^ yr^−1^, and are highly diverse in their provenance. We found that spatial carbon flows and local carbon fluxes are of the same order of magnitudes in freshwater and benthic ecosystems, suggesting an underlying dependency of these systems on resources provided by connected terrestrial and pelagic ecosystems respectively. By contrast, in terrestrial systems, cross-ecosystem subsidies were two to three orders of magnitude lower than local production (grasslands and forests), indicating a weaker quantitative influence on functioning. Those subsidies may still be qualitatively important, however, as some have high nutrient content^7,8^. We also find important gaps in carbon flow quantification, notably of cross-ecosystem subsidies driven by animal movements, which likely leads to general underestimations of the magnitude and direction of cross-ecosystem linkages^9^. Overall, we demonstrate strong ecosystem couplings, suggesting that ecosystems can be vulnerable to alterations of these flows and pointing to an urgent need to re-think ecosystem functioning in a spatial perspective.

## Main text

Ecosystems and the services that they provide are essential for material and cultural human welfare^10,11^, but paradoxically, human activities threaten ecosystem integrity^12,13^. Maintaining functional ecosystems, or restoring degraded ones, requires the identification of dominant mechanisms driving their dynamics. At the local scale, ecologists have accumulated extensive data on individual ecosystems’ functioning^14^. Accurate depictions of within-ecosystem fluxes, such as biomass and detritus production, respiration, or decomposition, exist for all broad ecosystems types, including terrestrial, freshwater and marine ones.

But ecosystems are not isolated. For flows of dispersing organisms the role on large-scale species coexistence and community dynamics is well studied^15,16^. However, it remains unclear to which extent ecosystem functioning also depends on cross-ecosystem flows of resources, such as in the form of detritus or nutrients^1,17^. According to the recently developed meta-ecosystem theory^16,18^, such cross-ecosystem resource flows can induce strong interdependencies between ecosystems and drive ecosystem functioning^3–6^. While the literature on subsidies provides emblematic cases of resources moving between ecosystems, including passive transport of leaves windblown from forests to streams^2^, or active transport such as aquatic insects emerging onto land^19^, we still lack a general quantification of these flows. The dispersion of data over many research areas and inconsistencies in the units of measurements used has hitherto precluded a general and synthetic overview of resource spatial flows.

Here, we conduct the first quantitative synthetic assessment of cross-ecosystem subsidies connecting the major ecosystem types across the globe (Figure 1). Specifically, we compare the magnitudes of cross-ecosystem subsidies to within-ecosystem fluxes in order to infer their relative contribution to ecosystem functioning. This also gives a basis to identify ecosystems’ vulnerability to increasing alterations of resource flows under the context of ongoing global changes^20^. We based our analysis on generally convertible estimates in units of carbon (gC m^2^ y^−1^; see Methods) and systematically searched for quantifications of spatial subsidies connecting terrestrial (forest, grassland, agroecosystem, desert), freshwater (stream, lake), and marine (pelagic and benthic) ecosystems. To compare these spatial flows, we also assembled comprehensive quantifications of local biological fluxes of carbon (i.e., gross primary production, ecosystem respiration, and decomposition; see Methods) within the different ecosystem types. We assembled 518 measurements of spatial flows and 2516 of local fluxes, reaching a total of 3034 data points extracted from 557 studies. Analysing this data set with its internally-consistent measurements in carbon units reveals for the first time the widespread importance of spatial resource flows to local ecosystem functioning.

**Figure 1.**
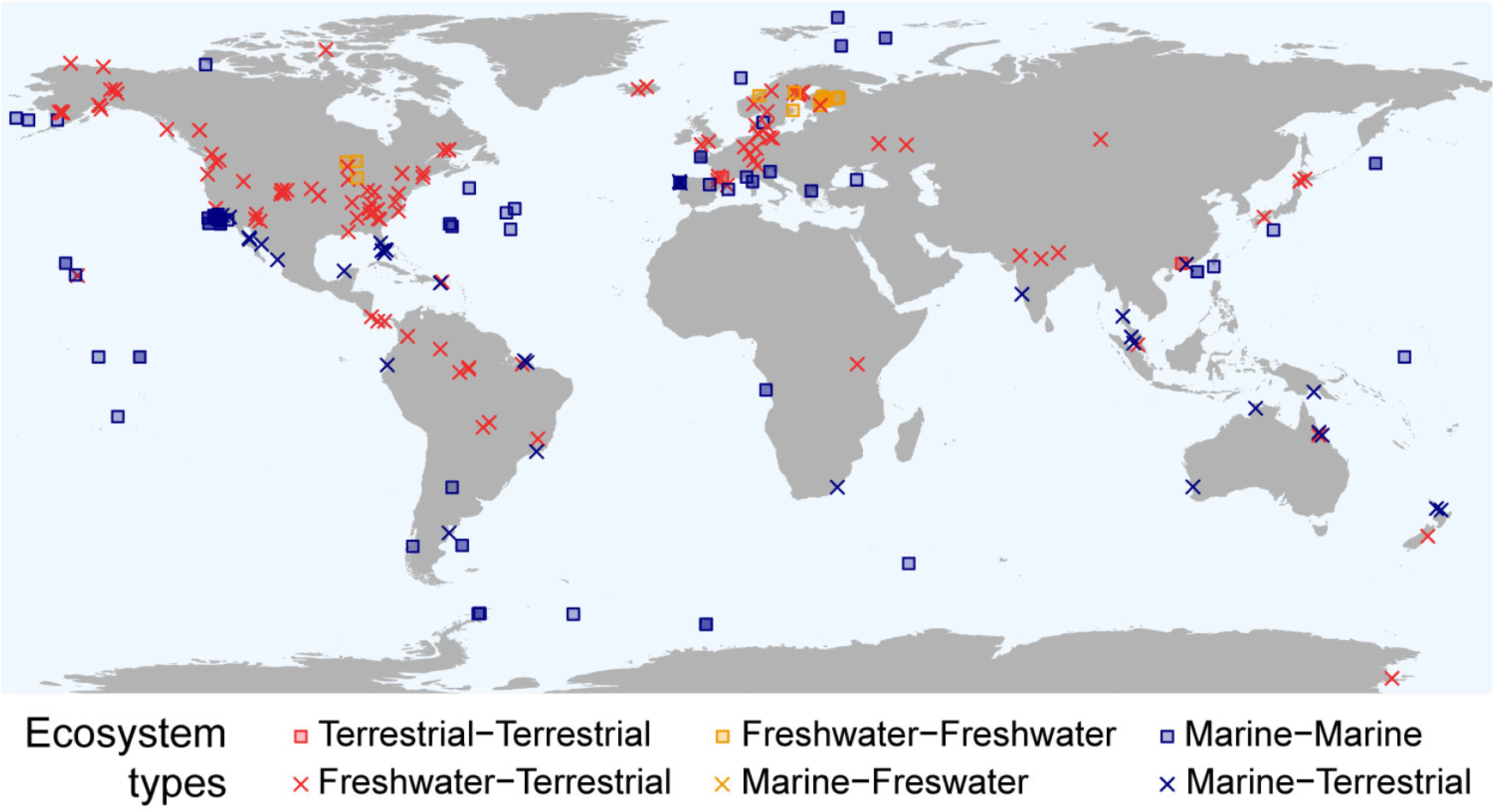
Global distribution of available cross-ecosystem subsidy data. Colours and shapes indicate the type of ecosystems coupled by cross-ecosystem subsidies: terrestrial (i.e., forest, grassland, agro-ecosystem, desert), marine (i.e., ocean pelagic, ocean benthic) and freshwater (i.e., stream, lake, wetland).

Spatial flows of cross-ecosystem carbon subsidies range over eight orders of magnitude of mass of carbon annually transported to ecosystems (gC m^−2^ y^−1^), from a few milligrams (aquatic insects deposited in forest) to more than ten tons per meter squared and year (wrack on shores) (Figure 2). The materials transported among ecosystems are as various as living animals, dead plants and animals, and dissolved carbon. Ecosystems occupying the lowest elevations in land-and seascapes receive downward flows of dead material or small organisms falling from above ecosystems, while terrestrial ecosystems receive more lateral flows, for instance of material transported by wind, ocean tides, or animal movements. Not surprisingly, ecosystems dominated by primary producer biomass (e.g., forests, grasslands, submarine meadows, and kelp forests), export the largest flows of primary production-derived material (median / interquartile range [IQR] of 148.7 / [47.1-246.6] and 164 / [93.2-1659.3] gC m^−2^ yr^−1^ for terrestrial vegetation and macro-algae, respectively). By comparison, invertebrate subsidies are substantially lower (median: 1.51 gC m^2^ y^−1^ / IQR: [0.44-5.41]), but might reach similar values in specific ecosystems (e.g., 124 gC m^2^ y^−1^ of aquatic insects flowing from lakes to tundras in Iceland). Lastly, few studies document spatial flows of vertebrate origin (3.1%), and those which do report highly contrasting values: faeces of foraging deer or fishery discards represent small flows of around 0.1-1 gC m^2^ y^−1^, while the net inputs to freshwater ecosystems of hippopotamus defecating into rivers, or of drowned migrating wildebeests, range between 100 and 1000 gC y^−1^ per meter squared of river. These disparities stress that the quantitative effect of carbon brought to ecosystems via large animals might greatly depend on how resource deposition is spatially constricted to small or large areas. Overall, our global picture highlights the ubiquity of cross-ecosystem subsidies, and the variety in their magnitude.

**Figure 2.**
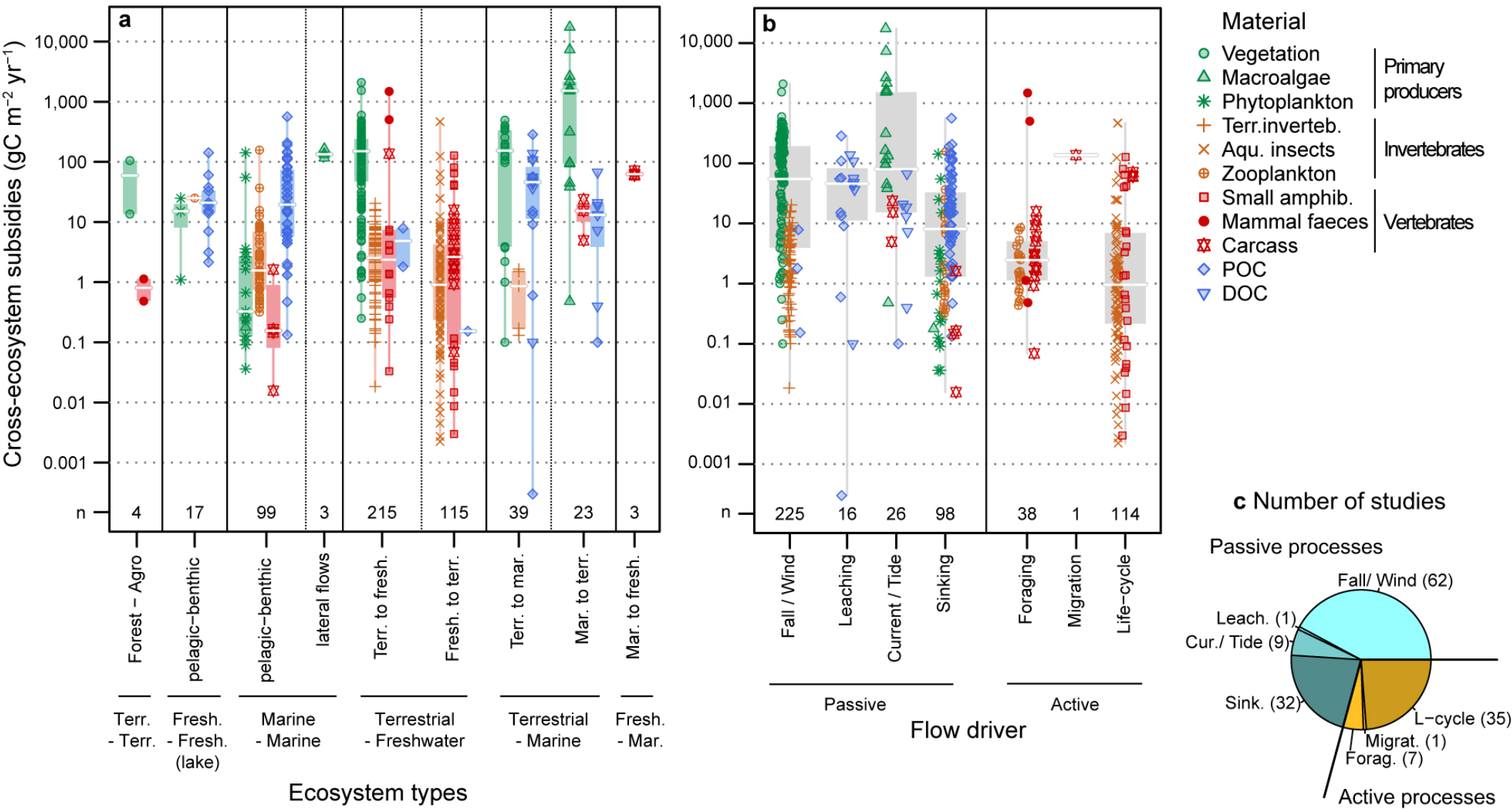
Architecture of cross-ecosystem carbon subsidies. Values are provided (**a**) by types of ecosystems connected by the subsidies, with vertical labels specifying ecosystems (“Agro” stands for “Agro-ecosystem”) or flow direction, and (**b**) by underlying flow driver, either via passive or active processes. Shapes and colours denote material transported. **a, b** - Bottom numbers, n, indicate the number of data points. Values are in gC m^−2^ y^−1^ on a log scale. The areal unit, m^2^, refers to the recipient ecosystem. Boxplots give median (white line), 25% and 75% percentiles (box), and range (whiskers). **c** - Number of studies by driver type.

To assess the importance of these subsidies to ecosystem functioning, we compared their magnitude to those of local biological fluxes within each receiving ecosystem. Magnitudes of subsidies versus local fluxes are similar in freshwater and in some benthic systems, whereas subsidies are generally far smaller than local fluxes in terrestrial ecosystems (Figure 3). This results both from more abundant cross-ecosystem subsidies in the direction of freshwater and benthic ecosystems than terrestrial and pelagic systems, and from substantially lower primary production in the former. Ecosystem respiration in freshwater and benthic systems often exceeds local primary production, despite a noticeable variability, for example due to differences in light availability in shallow tropical sea grass meadows *versus* deep waters promoting or constraining photosynthesis respectively. Ecosystem heterotrophy (see negative net ecosystem production in Extended data Figure 1) is associated with cross-ecosystem inflows of comparable or greater magnitude than local production, indicating that functioning in these ecosystems depends substantially on allochthonous resources. This dependency makes freshwater and deep unproductive benthic systems sensitive to alterations of these resources and, thus, to donor ecosystem dynamics^21,22^.

**Figure 3.**
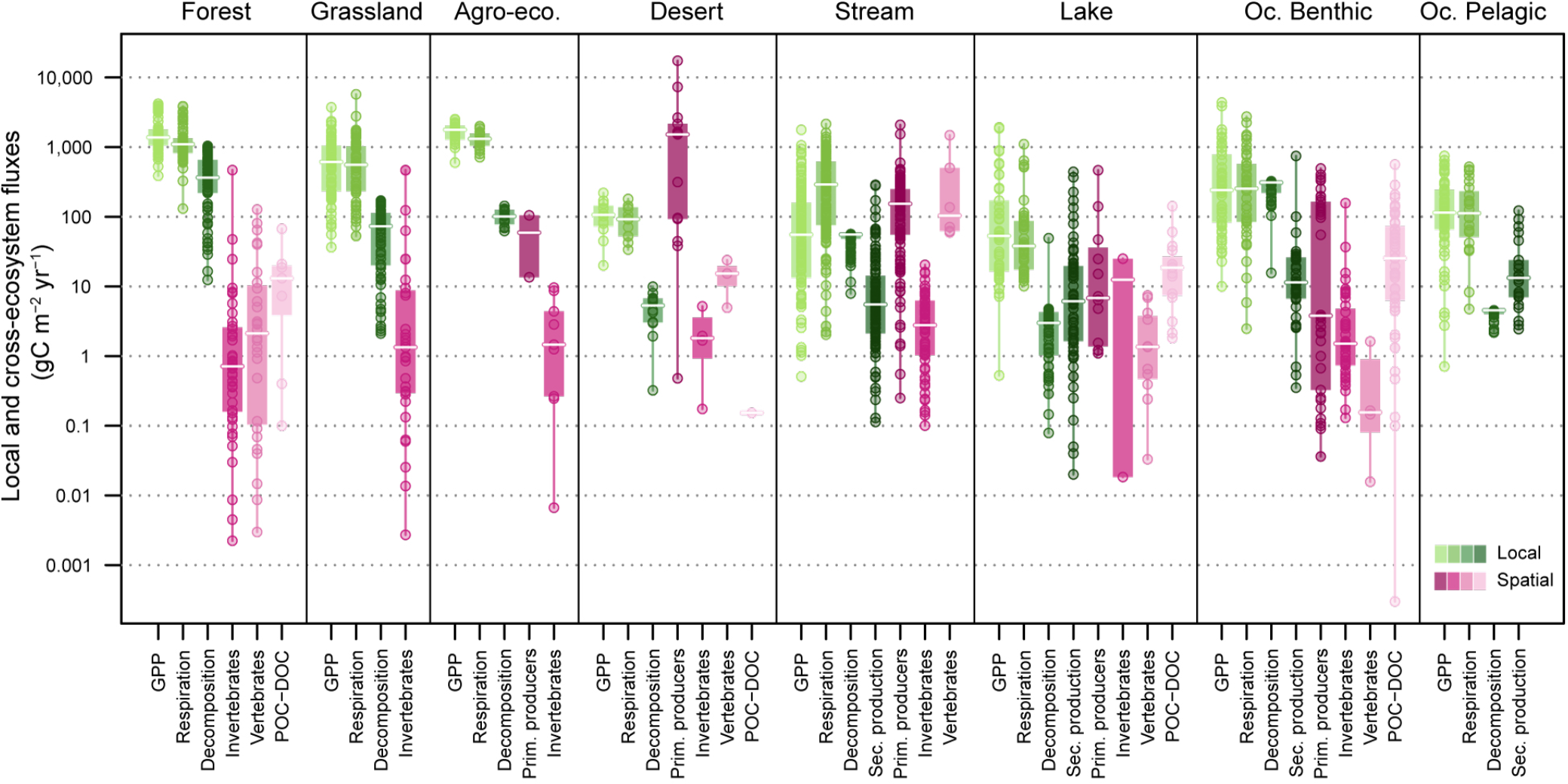
Comparison of local fluxes *versus* cross-ecosystem subsidies. Local fluxes within (green), and cross-ecosystem subsidies to (pink), specific ecosystem types (different panels). Represented local fluxes are gross primary production, ecosystem respiration, decomposition flux, and secondary production in aquatic systems. Cross-ecosystem subsidies are clumped into imported material of four origins: primary producers invertebrates, vertebrates, and organic carbon (particulate and dissolved). No subsidy to pelagic systems was documented. Circles give values in gC m^−2^ y^−1^ on a log scale. Boxplots give median (white line), 25% and 75% percentiles (box), and range (whiskers). Whisker and three null values of GPP in streams are omitted due to log scale.

By contrast, terrestrial and pelagic ecosystems tend to have a net autotrophic functioning and receive negligible subsidies compared to their local production, except in deserts (Figure 3). In the latter case, where local production is limited (e.g., by water availability), the potential impact of subsidies provided by other, less limited ecosystems (e.g., oceans) increases. Otherwise, autotrophic ecosystems seem relatively free from influence of spatial resources (or at least, from the influence of aquatic subsidies: we found only two studies reporting subsidies between terrestrial ecosystems). However, carbon subsidies are intimately linked to nutrients within biological molecules. Though quantitatively small in terms of carbon, subsidies to terrestrial systems are of higher quality with respect to nitrogen content than the vegetation-based subsidies they export (e.g. insects versus leaves in terrestrial-freshwater couplings^8,23^). Some cross-ecosystem subsidies, not considered here, are even predominantly of nitrogen or phosphorus, enriching terrestrial (e.g., guano from foraging sea birds^24^) or pelagic systems (e.g., excretion of marine mammals^25^). Carbon and nutrients in cross-ecosystem subsidies may relax different limitations in recipient ecosystems, thus enhancing regional production^6,26^.

Overall, our synthesis of carbon-based cross-ecosystem subsidies identifies strong spatial couplings between autotrophic and heterotrophic ecosystems (Figure 4). Cataloguing the types of subsidies documented so far, and their drivers (Figure 1), also reveals important gaps in the assessment of ecosystem couplings. A large portion of the quantified subsidies concerns flows occurring at terrestrial-freshwater (66.9%) or pelagic-benthic (18.0%) interfaces, and are driven by passive rather than active processes (Figure 2c; 70.5%). This may be because passive flows of detritus or of small organisms can be easily measured by various traps. By contrast, it is more challenging to quantify active spatial flows resulting from animal movement. This requires indirect estimations, for instance combining animal tracking and measurements of ingestion and excretion rates^27^. Quantifications are rare (Figure 2), but the few existing studies suggest that widespread animal movements, such as foraging or migration, could act as important ecosystem connectors^28^. In particular, mobile species using multiple ecosystems^29^, or those whose behaviour leads to massive aggregations of individuals, likely move important amounts of resources among ecosystems. At the moment, the lack of data hinders an assessment of the global importance of these actively-driven cross-ecosystem subsidies^9^. However, such linkages could render common ecosystem couplings more bi-directional than previously thought, as proposed for the aquatic-terrestrial interface^8,30^. Detailed evaluation of flow bi-directionality is essential to better understand spatial feedbacks between connected ecosystems. Resource exchanges between ecosystems underlie both optimization in resource use at the landscape scale, and potentials for spatial amplification of local perturbations.

**Figure 4.**
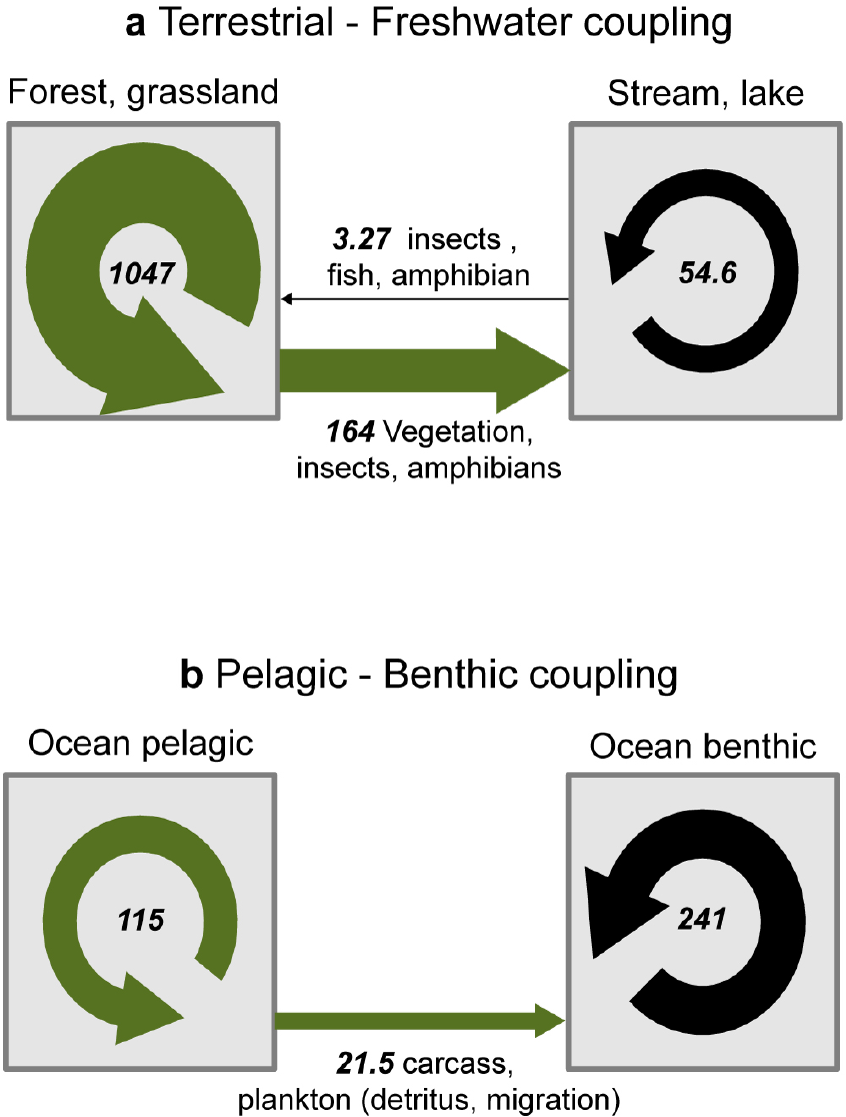
Well-documented natural meta-ecosystems. Cross-ecosystem subsidies suggest significant spatial couplings between (**a**) terrestrial and freshwater ecosystems and (**b**) pelagic and benthic areas in marine ecosystems. Round and horizontal arrows represent gross primary production (GPP) and cross-ecosystem subsidies, respectively. Numbers in italic are median values for GPP and sum of median values of cross-ecosystem subsidies of each different origin (primary producer, invertebrate, vertebrate, and particulate and dissolved organic carbon), both expressed in gC m^−2^ y^−1^. Width of arrows is proportional to these values (based on double squared root transformation).

In summary, the magnitude and importance of cross-ecosystem carbon subsidies for ecosystem functioning is globally variable, but predictable based on general knowledge of internal ecosystem fluxes and metabolism (net heterotroph vs. autotroph). Freshwater and benthic ecosystems might be especially vulnerable to subsidy alteration, and to perturbations in autotrophic ecosystems spatially cascading via these subsidies. However, gaps in subsidy quantification leave some uncertainty as to whether different ecosystems might act as buffers or amplifiers of spatial dynamics within landscapes. Documenting these ecological blind spots is necessary to improve our ability to predict ecosystem responses to global changes across landscapes.

## Methods

We conducted an extensive literature review of empirical values of cross-ecosystem subsidies over the globe (distribution in Figure 1), and compared their magnitude to local fluxes within ecosystems receiving these subsidies. We chose carbon as the focal material unit to profit from widely available carbon data for local fluxes. To enable the spatial flows to local flux comparison, we only considered measurements of spatial flows that were either provided in or could be converted into gC m^−2^ yr^−1^.

### Data collection

Our systematic search covered four broad categories of terrestrial ecosystems (forest, grassland, agro-ecosystem, and desert) and four of aquatic ecosystems (stream, lake, pelagic ocean and benthic ocean). We considered all ecosystems (if available) in five major global climatic zones (arctic/alpine, boreal, temperate, tropical and arid). Extended Data Table 1 provides the definitions of ecosystem categories and climatic zones. For marine ecosystems, we grouped arctic, boreal, temperate *versus* arid and tropical climates into “Cold” and “Warm” waters respectively, to account for a lesser influence of climate on oceanic systems due to the buffering effect of large water volumes. For each relevant ecosystem x climatic zone combination (see Extended Data Figure 2), we collected local carbon flux and spatial carbon flow data. We used all possible combinations of these categories and terms with similar meanings (see Extended Data Table 1) in our systematic search (see details in the next paragraphs).

We collected available values of subsidies linking the above-mentioned different ecosystems, and which could be converted into gC m^−2^ yr^−1^ in order to homogenize data and make comparisons possible. The latter constraint excluded cross-ecosystem flows of nutrients for which no carbon equivalent was possible, and flows expressed without information of the area of influence in the ecosystem receiving the subsidy. For instance, measurements of amount of dissolved organic carbon flowing from streams into estuaries^31^ where excluded because the impacted area was undefined. Terms primarily used for the search of spatial flows were “(subsid* OR spatial flow*) AND ecosystem”, with “ecosystem” also being replaced by specific ecosystems or pairwise combinations of the ecosystem types of interest.

In addition, for each ecosystem x climatic zone combination, we systematically searched published literature for values of the following within-ecosystem carbon fluxes: gross primary production (GPP), secondary production in aquatic ecosystems, ecosystem respiration (R_e_), net ecosystem production (NEP), and decomposition fluxes. Since decomposition fluxes were rarely directly provided, we derived them from detritus stocks and decomposition rates (see next section for calculations). A first systematic search was conducted by using all possible combinations of the names of each ecosystem type, climatic zone and flux of interest, with small variation when relevant (e.g. “decomposition OR decay” for decomposition flux and rates). The different terminologies used across various research fields to describe the same processes, and the fact that the data of interest were often located in different sections of the studies (Methods *versus* Results) limited the efficiency of standardized keyword search across the data types. We therefore complemented the dataset with multiple customized searches until we compiled a minimum number of ten independent values of each variable of interest (i.e. fluxes, detritus stock, and decomposition rate) for each ecosystem x climatic zone combination. At the end, data were pooled by ecosystem type.

In total, we collected 3,034 values from 557 published studies, including 518 values of cross-ecosystem subsidies. A summary of all values and the respective references are provided in Extended data Table 2 (cross-ecosystem subsidies) and Extended data Table 3 (local fluxes).

### Calculations used for data extraction

When only two of three major fluxes (production and respiration, and net ecosystem production (*GPP*, *R*_*e*_, and *NEP*, respectively) were reported, we estimated the unreported flux:

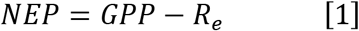

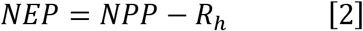

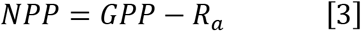

*NPP* is the net primary production, *R*_*h*_ the heterotrophic respiration and *R*_*a*_ the autotrophic respiration.

We derived decomposition fluxes *D*_*F*_ from detritus stocks *D*_*M*_ and decomposition rates *k*, with the classical exponential decay model:

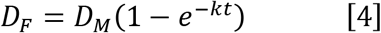

To calculate individual decomposition flux values, we parameterized detritus stock *D*_*M*_ with the median values of all detritus stocks in a given ecosystem x climatic zone combination and used the decomposition rate values collected from the literature to produce flux values. The rates were values of *k*, the first order constant in the classical exponential decay model. When not directly provided, we derived *k* with one of the equations proposed by Cebrian and Lartigue^14^ depending on the data available in the study:

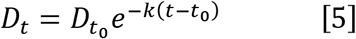

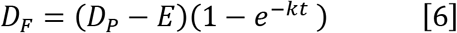

In Equation [5], *D*_*t*_ is the detrital mass at time *t* and *D*_*t_0_*_ the initial detrital mass. This equation was used when decomposition was estimated as the proportion of detrital mass loss (1 - *D*_*t*_/ *D*_*t*_0__) via a litter-bag experiment, a classical method in freshwater and terrestrial ecology. In equation [6], *D*_*F*_ is the (absolute) decomposition flux during the study period *t*, that is the flux from detritus stock to bacteria and other detritivores, *D*_*P*_ is the detritus production, and *E* the detritus export (e.g. sedimentation). In few cases of ocean pelagic data, we used the microbial loop of primary production *versus* bacterial production to parameterize *D*_*P*_ and *D*_*F*_, respectively. If not available, the export rate was set to 0, leading to *k* underestimation, which is conservative in our cross-ecosystem comparison given that *k* is already at the higher end of the range in these pelagic systems.

### Unit conversions

Once collected, we standardized values by converting them all into areal carbon units, that is, gC m^−2^ for detritus stocks and gC m^−2^ yr^−1^ for local fluxes and cross-ecosystem subsidies. Decomposition rates were expressed in yr^−1^.

#### Carbon conversion

We used data in carbon units (gC) when it was directly provided in the study, or we calculate the values using carbon content when reported in the study. Alternatively, we applied the most direct conversion of the data into carbon units depending on the level of detail available (see Extended data Table 4 for conversion factors). For decomposition rates, we did not transform units into carbon, assuming carbon loss rate to be identical to loss rate in the unit provided.

#### Time extrapolation

for local fluxes or rates provided in daily units, we extrapolated to the year by multiplying by 365 days in tropical systems where low or no seasonality is assumed. For the others, we multiplied by the number of days in the growing season as reported in the study, or the ice-free period in cold climates. When growing season length (GSL) was not specified in the study we used averaged estimates detailed by Garonna et al.^32^ for the different climatic zones in Europe^33^: 181 days for temperate climate (mean of atlantic and continental), 155 days for boreal, 116 days for arctic, and 163 days for arid systems (mean of Mediterranean and steppic). We did not apply any conversion if the value was measured on a study period longer than the above GSL for the corresponding climate.

#### Volume to area conversions and depth integration

Some data were given per unit of volume. For freshwater systems, we converted the data into area units by integrating them over the water column, using the mean depth of the river or lake. When not directly available in the study we calculated depth by dividing the volume per the area in lakes, or by estimating depth from discharge in rivers with the formula depth = *c.Q*^*f*^, with *c* = 0.2, *f* = 0.4 and *Q* the discharge in m^3^ s^−1^ ^34^. For small catchment areas, that is <1 km^2^, we estimated the depth to be 5 cm based on known river scaling-properties^34^. For marine data, notably production in the pelagic zone, studies generally provide a meaningful depth, which defines the euphotic zone such as the Secchi depth or the 1% light inflow depth. We integrated values in volume units over this depth, and to 100 m depth when only sampling depths were provided.

#### Areal units for cross-ecosystem subsidies

Since we were interested in comparing the magnitude of spatial subsidies to that of the recipient ecosystem’s own local fluxes, we needed cross-ecosystem subsidies measured in areal units of the recipient ecosystem. Depending on the method of quantification, cross-ecosystem flow can be directly expressed in this way (e.g. litter traps in the recipient ecosystem). In other cases, for example with vertical fluxes between pelagic and benthic ecosystems, the equivalence of donor and recipient ecosystem areas affected by the flow is obvious. However, in the case of lateral flows occurring from aquatic to terrestrial ecosystems (e.g. emergent insects, carcasses of salmon caught by bears), flows were often provided per m^2^ of donor area. We could not use such measurements directly because the magnitude of the flow in the recipient ecosystem depends on both the total surface of production and the boundary length. For instance, lakes with the same area but having circular *versus* complex-shorelines will lead, for the same total emergent insect flux, to higher *versus* lower magnitudes of flows, respectively, distributed per areal unit of the recipient ecosystem. Moreover, the maximum influence of such cross-ecosystem flow is found near at the shoreline and decreases with distance from the shore (for example, in aquatic insects^19^). To adjust for this in a conservative approach, we homogenized our aquatic-to-terrestrial flows assuming a uniform distribution of the flow on the first ten meters from the shore, a distance within which most of the aquatic insect flows fall^35^ (but could fall substantially farther in some specific systems^36^), or most of the salmon carcasses brought by bears on land are deposited^37^. Therefore, when the values of spatial flows were provided in areal unit of the donor ecosystem (aquatic), we first calculated the flow per meter shoreline (in some cases, values were already provided per meter shoreline; e.g., wrack deposited on beaches), and then divided this value by 10 meters of influenced land. To calculate the spatial flow per shoreline length in freshwater systems, we followed the method described by Gratton et al.^19^ for insect emergence data: In lakes we multiplied the data per the total lake area and divided per the perimeter. When not directly available, the perimeter was approximated by 2*D_L_*(area*π)^1/2^, with D_L_ the development factor defined by Kalff^38^, which is 1 for circular shapes, 2 for the same area with a twofold larger perimeter. In streams we multiplied the original donor area flow value by stream width, and then divided it per two (riversides) to obtain the flow per shoreline length. The Extended Data Figure S3 show that our conclusions are not sensitive to the choice of the distance from shoreline used to calculate the recipient terrestrial area of aquatic subsidies: using a more conservative threshold of 100m only decreases the importance of spatial subsidies for terrestrial ecosystems, that we already assess as low.

### Flow drivers

We defined categories of flow drivers to examine the underlying processes of documented cross-ecosystem subsidies. Drivers could be either passive, via physical processes such as gravity, wind, water currents, tides, diffusion, or active, via animal movements such as those triggered by foraging behaviours, seasonal migration or cross-ecosystem movement of animals needed to complete a life-cycle. We kept these last three categories for active drivers and we clumped the passive drivers into categories reflecting broad classes of spatial flows: “Fall/wind” for aerial transport from terrestrial systems, “leaching” for diffusion processes, “current, tides” for lateral flows from aquatic systems and “sinking” for vertical passive flows in the water column (see Extended data Table 5). We analysed the data and plotted the figures with the software R^39^ and the R-packages ggmap^40^, maps^41^, and pgirmess^42^.

## Data availability

Data that support the findings of this study are summarized in Extended Data Table 2 and Extended Data Table 3, with all associated references in Supplementary Table 1.

## Acknowledgements

We thank Marcel Holyoak and Emanuel A. Fronhofer for discussions, and Mary O’Connor for comments on the manuscript.

## Author contribution

I.G., C.J.L., E.H. and F.A. conceived the study and collected the data. I.G. analysed the data, produced the figures and wrote the first draft of the manuscript. All authors edited and contributed to the final version of the manuscript.

## Extended Data Display

**Extended Data Figure 1.**
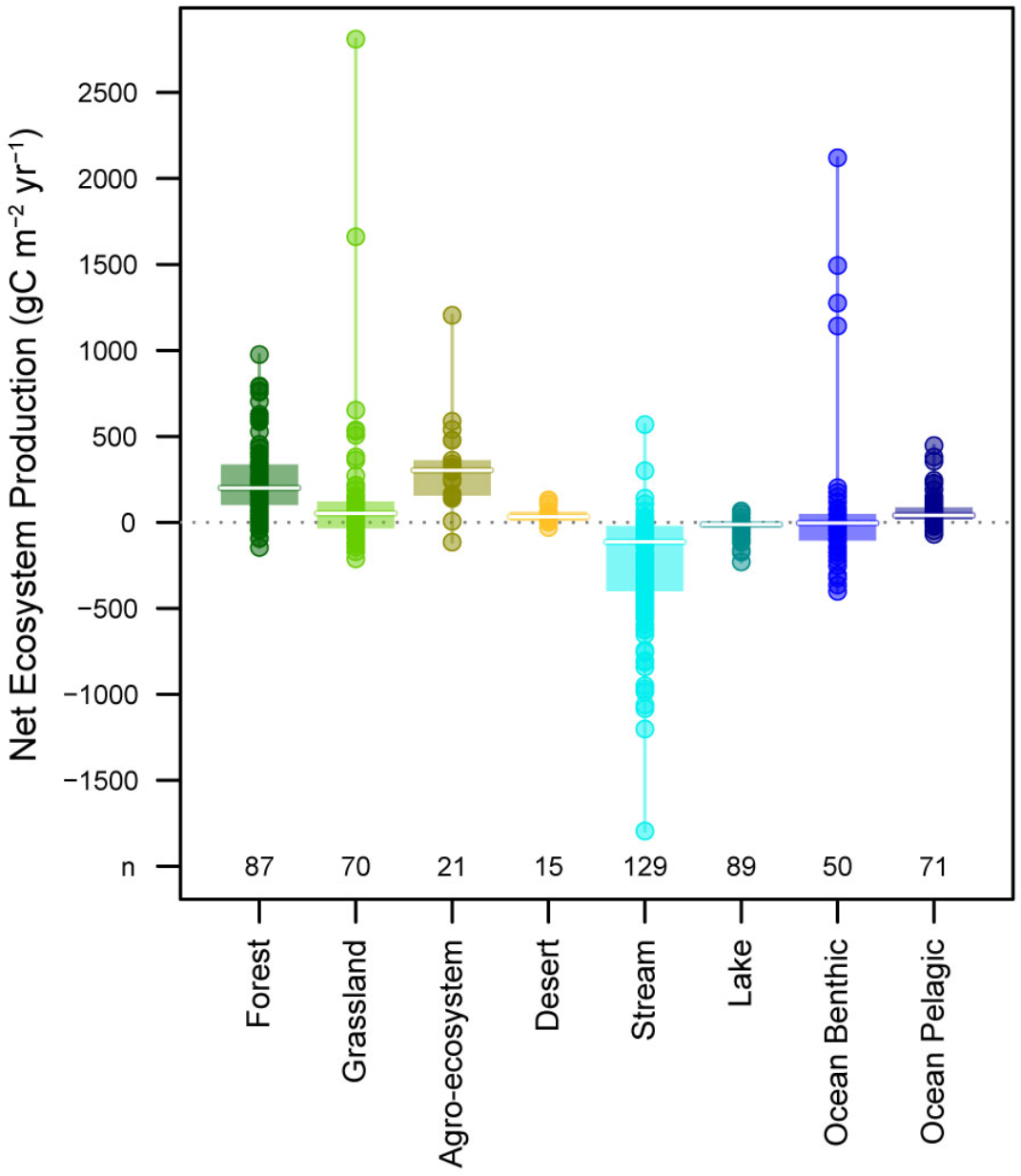
Net Ecosystem Production for different ecosystem types. Net ecosystem production corresponds to the balance between gross primary production and ecosystem respiration. Negative values denote net heterotrophic functioning. Circles give individual values in gC m^−2^ y^−1^. Note that these are production fluxes and not productivity rates (biomass turnover), for which we would have higher values in aquatic compared to terrestrial systems. Boxplots give median (white line), 25% and 75% percentiles (box), and range (whiskers). Bottom numbers (n) indicate the number of data points.

**Extended Data Figure 2.**
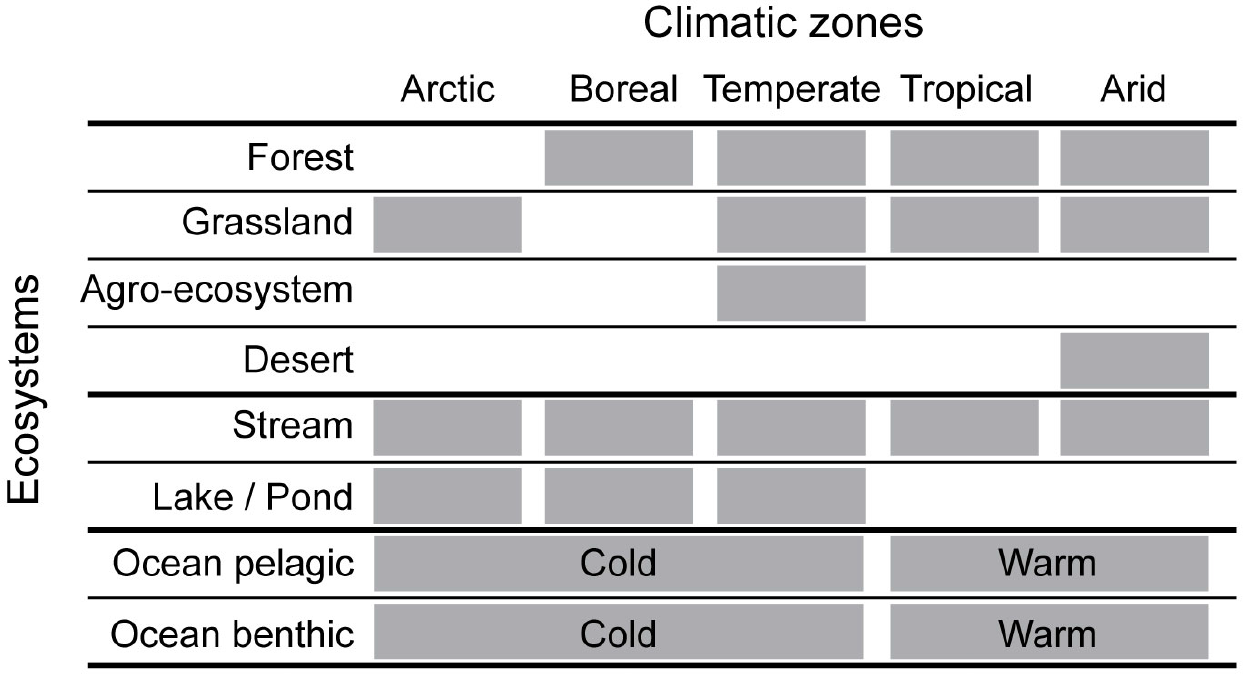
Data collection. Combinations of ecosystem types and climatic zones considered in this study. We collected values of local carbon fluxes for all combinations marked by a grey rectangle, as well as cross-ecosystem subsidies linking these ecosystems. The other combinations either do not exist, or do not have enough data available.

**Extended Data Figure 3.**
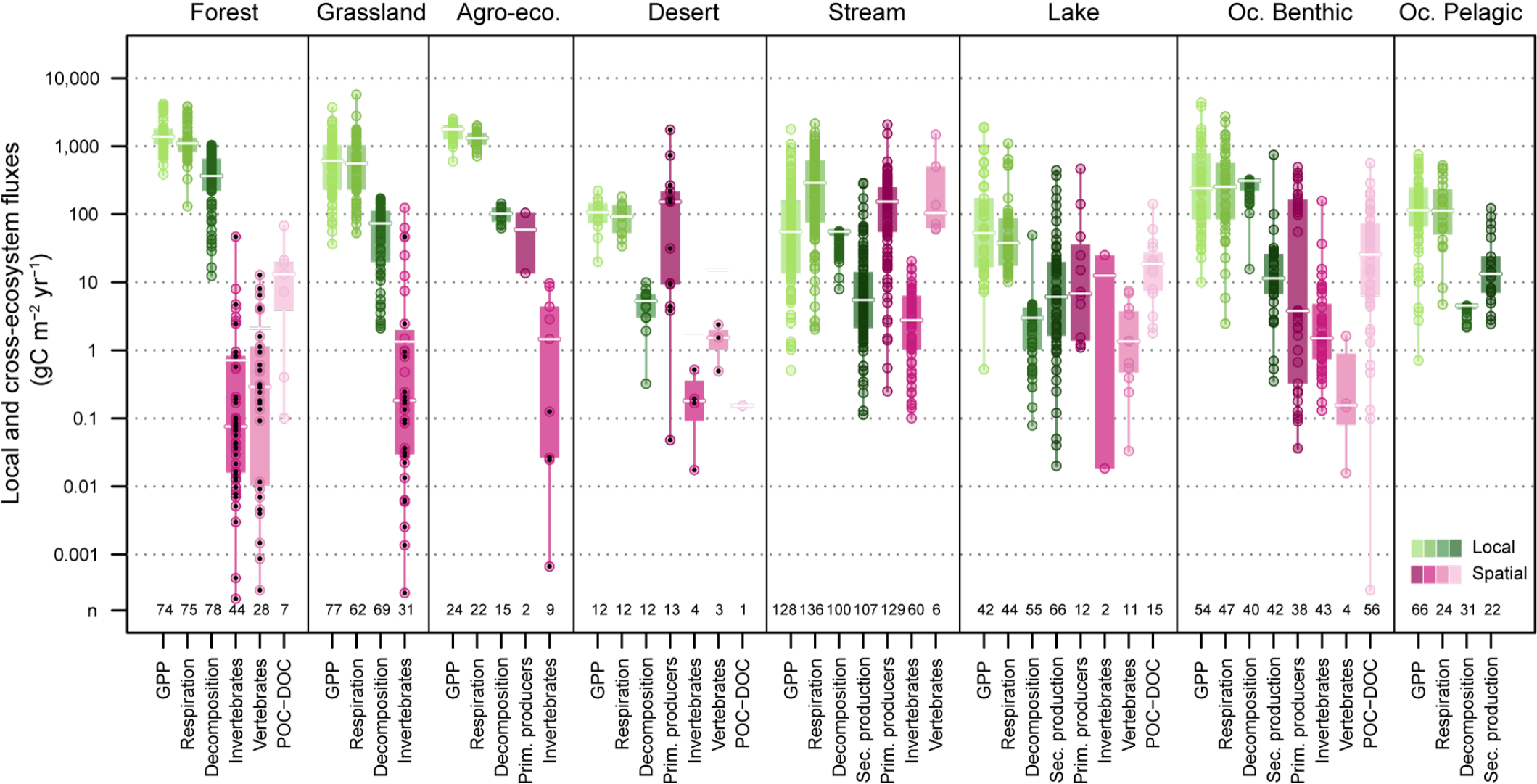
Sensitivity of local fluxes *versus* cross-ecosystem subsidies to shoreline threshold. Local fluxes within-(green) and cross-ecosystem subsidies to-(pink), specific ecosystem types (different panels). A threshold of 100 meters from shoreline (rather than 10m in Figure 3) was used to calculate the recipient terrestrial area influenced by aquatic cross-ecosystem subsidies. Circles give values in gC m^−2^ y^−1^ on a log scale. Black dots show the data points influenced by this choice. Boxplots give median (white line), 25% and 75% percentiles (box), and range (whiskers). Whisker and three null values of GPP in streams are omitted due to log scale. Bottom numbers (n) indicate the number of data points.

**Extended Data Table 1.**
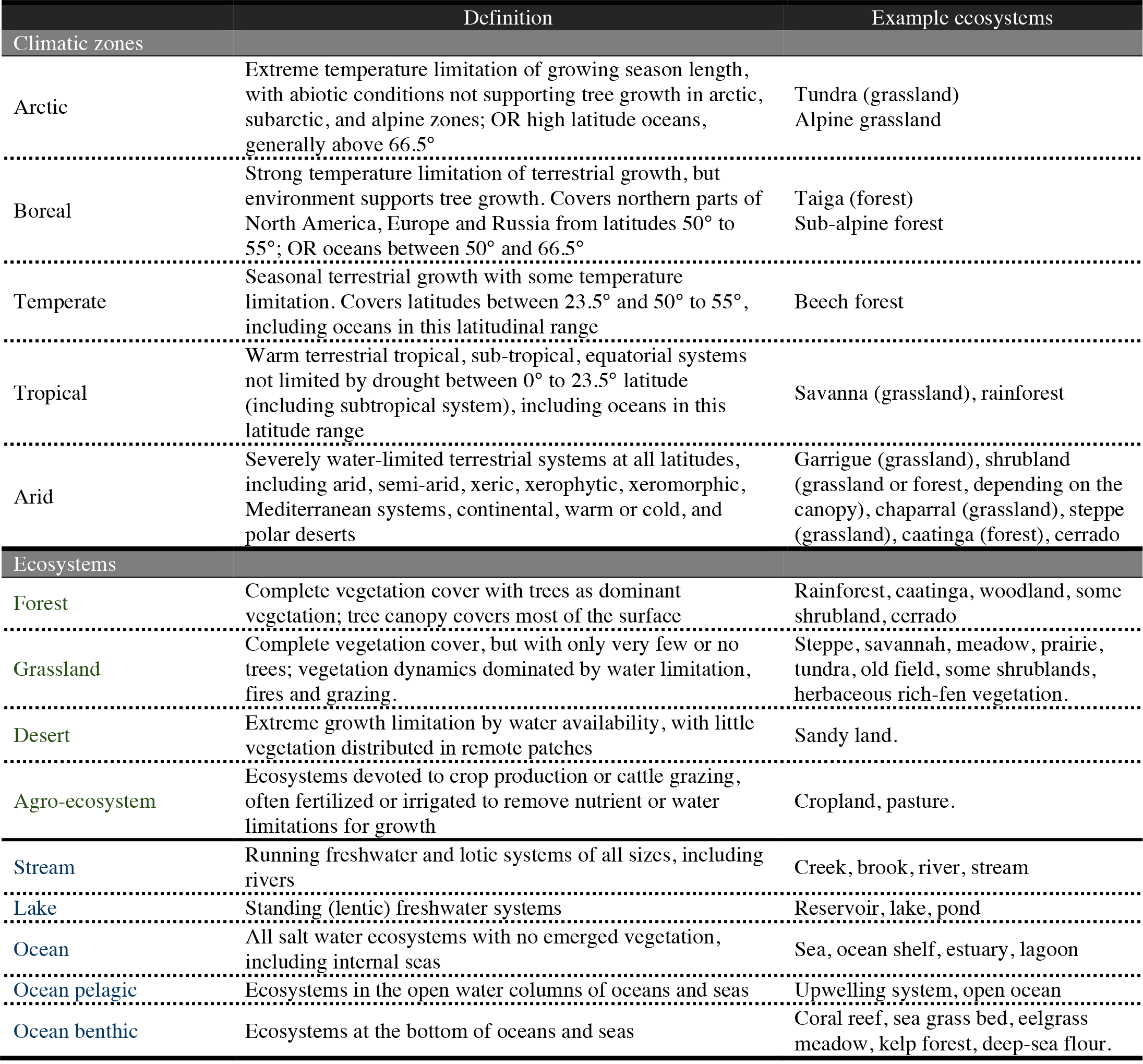
Definition of ecosystem and climate categories.

|  | Definition | Example ecosystems |
| --- | --- | --- |
| **Climatic zones** |  |  |
| Arctic | Extreme temperature limitation of growing season length, with abiotic conditions not supporting tree growth in arctic, subarctic, and alpine zones; OR high latitude oceans, generally above 66.5° | Tundra (grassland)<br>Alpine grassland |
| Boreal | Strong temperature limitation of terrestrial growth, but environment supports tree growth. Covers northern parts of North America, Europe and Russia from latitudes 50° to 55°; OR oceans between 50° and 66.5° | Taiga (forest)<br>Sub-alpine forest |
| Temperate | Seasonal terrestrial growth with some temperature limitation. Covers latitudes between 23.5° and 50° to 55°, including oceans in this latitudinal range | Beech forest |
| Tropical | Warm terrestrial tropical, sub-tropical, equatorial systems not limited by drought between 0° to 23.5° latitude (including subtropical system), including oceans in this latitude range | Savanna (grassland), rainforest |
| Arid | Severely water-limited terrestrial systems at all latitudes, including arid, semi-arid, xeric, xerophytic, xeromorphic, Mediterranean systems, continental, warm or cold, and polar deserts | Garrigue (grassland), shrubland (grassland or forest, depending on the canopy), chaparral (grassland), steppe (grassland), caatinga (forest), cerrado |
| **Ecosystems** |  |  |
| Forest | Complete vegetation cover with trees as dominant vegetation; tree canopy covers most of the surface | Rainforest, caatinga, woodland, some shrubland, cerrado |
| Grassland | Complete vegetation cover, but with only very few or no trees; vegetation dynamics dominated by water limitation, fires and grazing. | Steppe, savannah, meadow, prairie, tundra, old field, some shrublands, herbaceous rich-fen vegetation. |
| Desert | Extreme growth limitation by water availability, with little vegetation distributed in remote patches | Sandy land. |
| Agro-ecosystem | Ecosystems devoted to crop production or cattle grazing, often fertilized or irrigated to remove nutrient or water limitations for growth | Cropland, pasture. |
| Stream | Running freshwater and lotic systems of all sizes, including rivers | Creek, brook, river, stream |
| Lake | Standing (lentic) freshwater systems | Reservoir, lake, pond |
| Ocean | All salt water ecosystems with no emerged vegetation, including internal seas | Sea, ocean shelf, estuary, lagoon |
| Ocean pelagic | Ecosystems in the open water columns of oceans and seas | Upwelling system, open ocean |
| Ocean benthic | Ecosystems at the bottom of oceans and seas | Coral reef, sea grass bed, eelgrass meadow, kelp forest, deep-sea floor. |

**Extended Data Table 2.** Summary of cross-ecosystem subsidy data.

| Donor | Recipient | Material | n | Min | q25 | Median | q75 | Max | Data References |
| --- | --- | --- | --- | --- | --- | --- | --- | --- | --- |
| Forest | Agroeco. | Vegetation | 2 | 13.545 | 36.360 | 59.175 | 81.990 | 104.81 | 50 |
| Forest | Stream | Vegetation | 96 | 0.2500 | 55.247 | 178.65 | 280.62 | 2,085.8 | 8, 12, 13, 14, 15, 16, 17, 21, 26, 27, 28, 29, 30, 31, 33, 65, 78, 90, 96, 117, 118, 123, 124, 138, 203, 205, 212, 235, 375, 389, 392 |
| Forest | Stream | Terr.inverteb. | 49 | 0.1414 | 1.0683 | 3.0368 | 7.3321 | 20.386 | 2, 3, 4, 5, 6, 7, 23, 25, 32, 65, 98, 147, 365, 369, 370, 375, 525, 526 |
| Forest | Lake | Vegetation | 9 | 1.2000 | 1.5439 | 6.4000 | 47.000 | 465.00 | 20, 109, 295, 316, 375, 520, 521 |
| Forest | Lake | Terr.inverteb. | 1 | 0.0184 | 0.0184 | 0.0184 | 0.0184 | 0.0184 | 375 |
| Forest | Lake | Small amphib. | 11 | 0.0331 | 0.4706 | 1.3528 | 3.7450 | 7.4100 | 60, 61 |
| Forest | Lake | POC | 2 | 1.8100 | 3.3259 | 4.8418 | 6.3576 | 7.8735 | 522 |
| Forest | Ocean | Vegetation | 18 | 0.1000 | 27.249 | 153.80 | 312.00 | 492.80 | 430, 523 |
| Forest | Ocean | Terr.inverteb. | 5 | 0.1300 | 0.1700 | 0.8600 | 1.5100 | 1.6400 | 527 |
| Forest | Ocean | POC | 9 | 0.0003 | 9.1000 | 15.100 | 56.900 | 285.00 | 430 |
| Forest | Ocean | DOC | 7 | 0.1000 | 40.500 | 48.000 | 82.050 | 138.10 | 430 |
| Grassland | Stream | Vegetation | 9 | 0.5500 | 16.000 | 64.593 | 250.00 | 265.77 | 16, 18, 65, 89, 117, 118, 138, 205 |
| Grassland | Stream | Terr.inverteb. | 6 | 0.1005 | 0.3413 | 0.8304 | 2.0334 | 2.5296 | 9, 32, 365 |
| Grassland | Stream | Mammal faeces | 2 | 502.00 | 746.75 | 991.50 | 1,236.2 | 1,481.0 | 364 |
| Grassland | Stream | Carcass | 1 | 136.80 | 136.80 | 136.80 | 136.80 | 136.80 | 556 |
| Agroeco. | Forest | Mammal faeces | 2 | 0.4829 | 0.6448 | 0.8067 | 0.9686 | 1.1305 | 367 |
| Agroeco. | Stream | Vegetation | 23 | 1.4000 | 61.814 | 104.70 | 165.15 | 264.80 | 12, 78 |
| Agroeco. | Stream | Terr.inverteb. | 5 | 1.4544 | 4.4640 | 5.2699 | 6.1504 | 6.1504 | 3, 98, 365 |
| Desert | Stream | Vegetation | 1 | 8.4000 | 8.4000 | 8.4000 | 8.4000 | 8.4000 | 94 |
| Wetland | Forest | Small amphib. | 1 | 64.634 | 64.634 | 64.634 | 64.634 | 64.634 | 59 |
| Wetland | Grassland | Aqu. insects | 6 | 0.0136 | 0.0339 | 0.1410 | 0.3297 | 2.4408 | 57, 66 |
| Stream | Forest | Aqu. insects | 30 | 0.0022 | 0.1377 | 0.4857 | 0.9384 | 8.3480 | 10, 11, 19, 65, 67, 74, 77, 78, 147, 266, 365, 366, 477 |
| Stream | Forest | Carcass | 14 | 0.0693 | 1.7076 | 2.9078 | 5.8411 | 15.980 | 386, 390 |
| Stream | Grassland | Aqu. insects | 5 | 0.0027 | 0.0628 | 0.1329 | 0.2828 | 0.9516 | 65, 77, 365 |
| Stream | Agroeco. | Aqu. insects | 9 | 0.0067 | 0.2648 | 1.4607 | 4.3822 | 9.6100 | 69, 77, 78, 365 |
| Stream | Desert | Aqu. insects | 4 | 0.1742 | 1.3000 | 1.8098 | 2.7536 | 5.1815 | 22, 34, 387 |
| Lake | Forest | Aqu. insects | 14 | 0.0976 | 0.7246 | 1.8110 | 9.1252 | 467.87 | 62, 368, 374, 518, 524, 528 |
| Lake | Forest | Small amphib. | 11 | 0.0030 | 0.0273 | 0.0914 | 41.002 | 127.33 | 58, 60, 528 |
| Lake | Grassland | Aqu. insects | 20 | 0.3057 | 1.2527 | 4.7484 | 24.641 | 467.87 | 1, 24, 62, 368, 374, 383, 517, 519 |
| Lake | Desert | POC | 1 | 0.1530 | 0.1530 | 0.1530 | 0.1530 | 0.1530 | 56 |
| L. Pelagic | L. Benthic | Phytoplankton | 3 | 1.0971 | 8.1016 | 15.106 | 19.965 | 24.824 | 99, 134, 379 |
| L. Pelagic | L. Benthic | Zooplankton | 1 | 24.983 | 24.983 | 24.983 | 24.983 | 24.983 | 379 |
| L. Pelagic | L. Benthic | POC | 13 | 2.1448 | 13.950 | 20.925 | 32.860 | 141.82 | 294, 296, 485 |
| Ocean | Forest | POC | 1 | 0.1000 | 0.1000 | 0.1000 | 0.1000 | 0.1000 | 430 |
| Ocean | Forest | DOC | 6 | 0.4000 | 8.7500 | 15.900 | 20.350 | 67.300 | 430 |
| Ocean | Desert | Macroalgae | 6 | 44.520 | 93.921 | 835.79 | 1,638.3 | 2,646.9 | 103, 104, 371 |
| Ocean | Desert | Carcass | 3 | 4.9500 | 10.125 | 15.300 | 19.575 | 23.850 | 104 |
| Ocean | Stream | Carcass | 3 | 59.667 | 61.075 | 62.483 | 66.847 | 71.211 | 389 |
| Oc. Pelagic | Oc. Benthic | Macroalgae | 1 | 0.1786 | 0.1786 | 0.1786 | 0.1786 | 0.1786 | 503 |
| Oc. Pelagic | Oc. Benthic | Phytoplankton | 16 | 0.0362 | 0.1222 | 0.4978 | 2.7648 | 142.00 | 310, 311, 499, 539 |
| Oc. Pelagic | Oc. Benthic | Zooplankton | 38 | 0.3167 | 0.7563 | 1.5629 | 6.4680 | 156.90 | 308, 311, 384, 385, 495, 539, 557 |
| Oc. Pelagic | Oc. Benthic | Carcass | 4 | 0.0157 | 0.1134 | 0.1556 | 0.5290 | 1.6200 | 499, 500, 503, 504 |
| Oc. Pelagic | Oc. Benthic | POC | 40 | 0.1330 | 5.7875 | 19.444 | 70.250 | 562.91 | 80, 102, 116, 131, 133, 304, 306, 311, 319, 330, 334, 335, 338, 393, 494, 514 |
| Oc. Benthic | Desert | Macroalgae | 7 | 0.4795 | 176.46 | 1,522.1 | 4,730.8 | 17,365 | 372, 373, 382 |
| Oc. Benthic | Oc. Benthic | Macroalgae | 3 | 115.80 | 124.65 | 133.50 | 148.75 | 164.00 | 418, 420 |

Agroeco. = Agro-ecosystem; Oc. = Ocean; Terr. = Terrestrial; Aqu. = Aquatic; inverteb. = invertebrates;
amphib. = amphibians; POC = Particulate Organic Carbon; DOC = Dissolved Organic Carbon.
We provide the number of data points (n), the minimum, 25% quartile (q25), median, 75% quartile (q75) and maximum values of estimates distribution for each pair of ecosystems coupled and per type of the material exchanged, as well as the associated references from which we extracted the data, listed in

**Extended Data Table 3.**
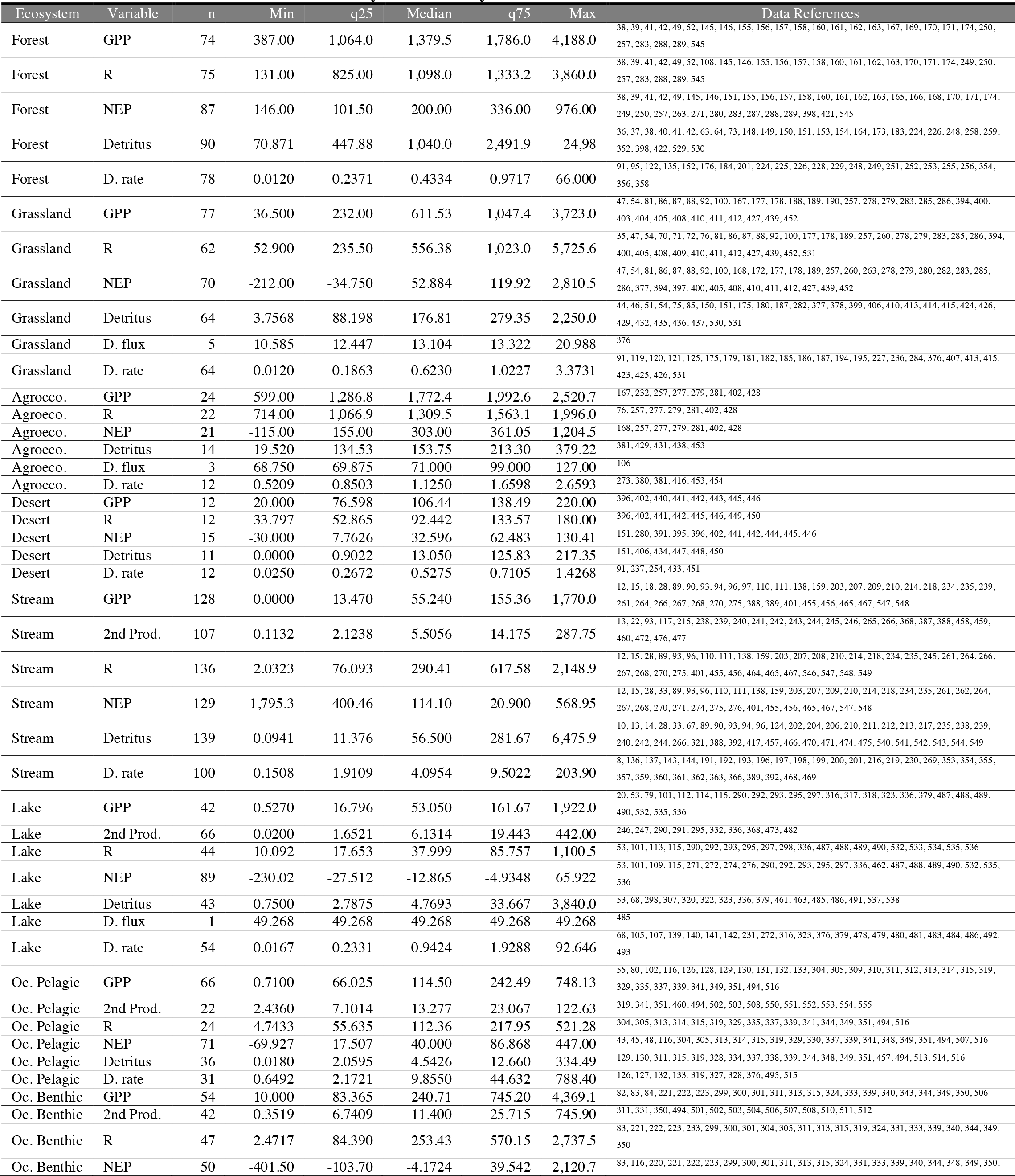
Summary of local ecosystem data.

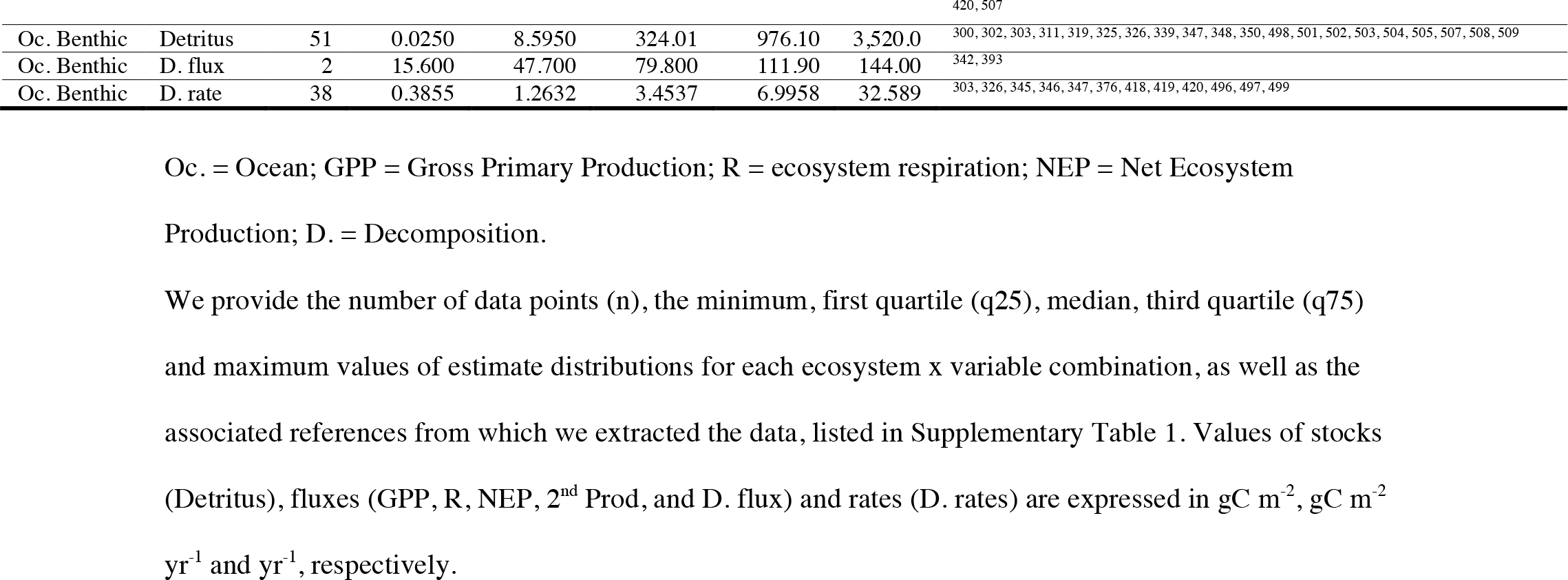

**Extended Data Table 4.**
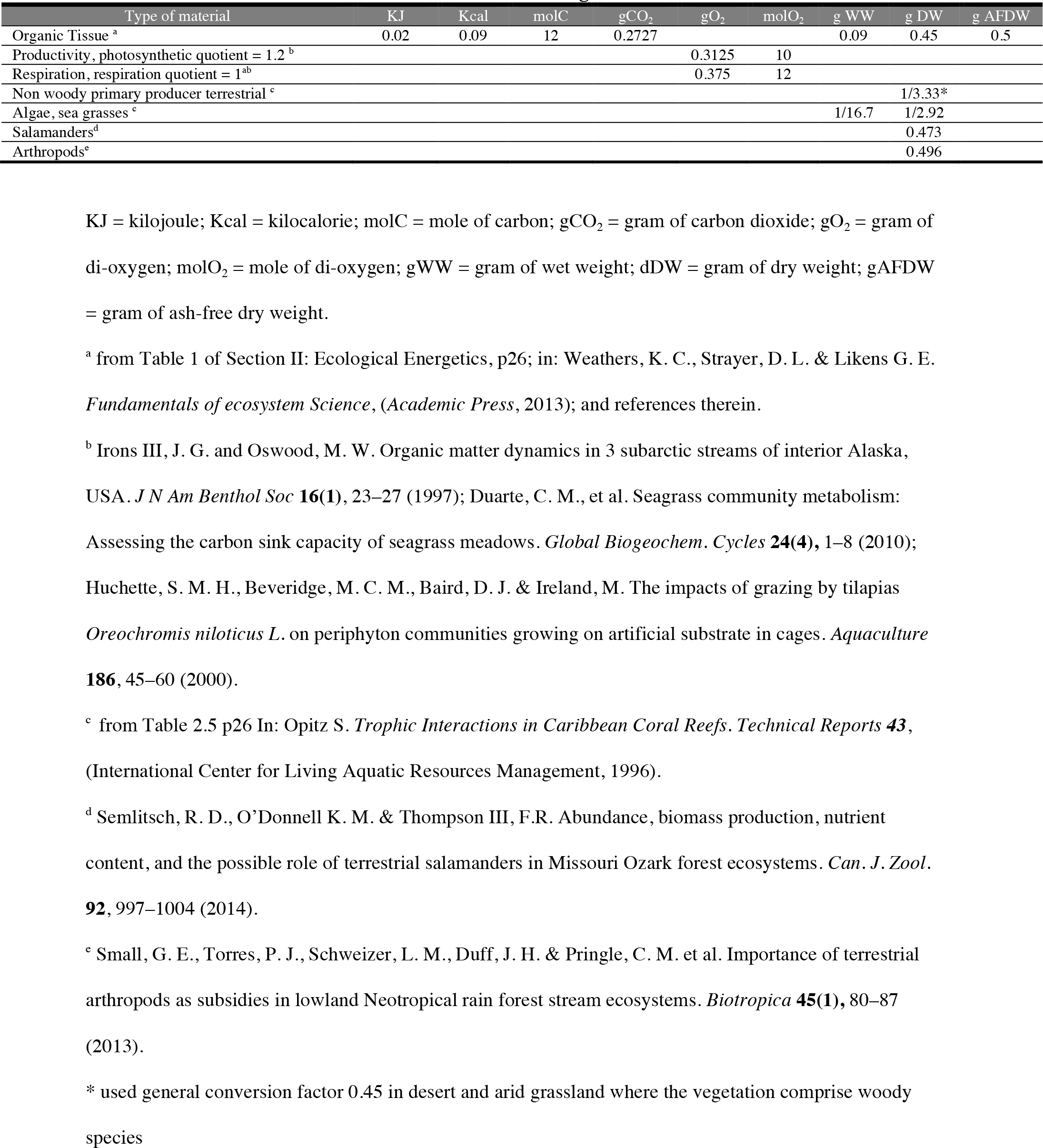
Factors used for conversions into grams of carbon.

| Type of material | KJ | Kcal | molC | gCO<sub>2</sub> | gO<sub>2</sub> | molO<sub>2</sub> | g WW | g DW | g AFDW |
| --- | --- | --- | --- | --- | --- | --- | --- | --- | --- |
| Organic Tissue <sup>a</sup> | 0.02 | 0.09 | 12 | 0.2727 |  |  | 0.09 | 0.45 | 0.5 |
| Productivity, photosynthetic quotient = 1.2 <sup>b</sup> |  |  |  |  | 0.3125 | 10 |  |  |  |
| Respiration, respiration quotient = 1 <sup>ab</sup> |  |  |  |  | 0.375 | 12 |  |  |  |
| Non woody primary producer terrestrial <sup>c</sup> |  |  |  |  |  |  |  | 1/3.33* |  |
| Algae, sea grasses <sup>c</sup> |  |  |  |  |  |  | 1/16.7 | 1/2.92 |  |
| Salamanders<sup>d</sup> |  |  |  |  |  |  |  | 0.473 |  |
| Arthropods<sup>e</sup> |  |  |  |  |  |  |  | 0.496 |  |

KJ = kilojoule; Kcal = kilocalorie; molC = mole of carbon; gCO<sub>2</sub> = gram of carbon dioxide; gO<sub>2</sub> = gram of di-oxygen; molO<sub>2</sub> = mole of di-oxygen; gWW = gram of wet weight; dDW = gram of dry weight; gAFDW = gram of ash-free dry weight.
<sup>a</sup> from Table 1 of Section II: Ecological Energetics, p26; in: Weathers, K. C., Strayer, D. L. & Likens G. E. *Fundamentals of ecosystem Science*, (Academic Press, 2013); and references therein.
<sup>b</sup> Irons III, J. G. and Oswood, M. W. Organic matter dynamics in 3 subarctic streams of interior Alaska, USA. *J N Am Benthol Soc* **16**(1), 23–27 (1997); Duarte, C. M., et al. Seagrass community metabolism: Assessing the carbon sink capacity of seagrass meadows. *Global Biogeochem. Cycles* **24**(4), 1–8 (2010); Huchette, S. M. H., Beveridge, M. C. M., Baird, D. J. & Ireland, M. The impacts of grazing by tilapias *Oreochromis niloticus* L. on periphyton communities growing on artificial substrate in cages. *Aquaculture* **186**, 45–60 (2000).
<sup>c</sup> from Table 2.5 p26 In: Opitz S. *Trophic Interactions in Caribbean Coral Reefs. Technical Reports* **43**, (International Center for Living Aquatic Resources Management, 1996).
<sup>d</sup> Semlitsch, R. D., O'Donnell K. M. & Thompson III, F.R. Abundance, biomass production, nutrient content, and the possible role of terrestrial salamanders in Missouri Ozark forest ecosystems. *Can. J. Zool.* **92**, 997–1004 (2014).
<sup>e</sup> Small, G. E., Torres, P. J., Schweizer, L. M., Duff, J. H. & Pringle, C. M. et al. Importance of terrestrial arthropods as subsidies in lowland Neotropical rain forest stream ecosystems. *Biotropica* **45**(1), 80–87 (2013).
\* used general conversion factor 0.45 in desert and arid grassland where the vegetation comprise woody species

**Extended Data Table 5.**
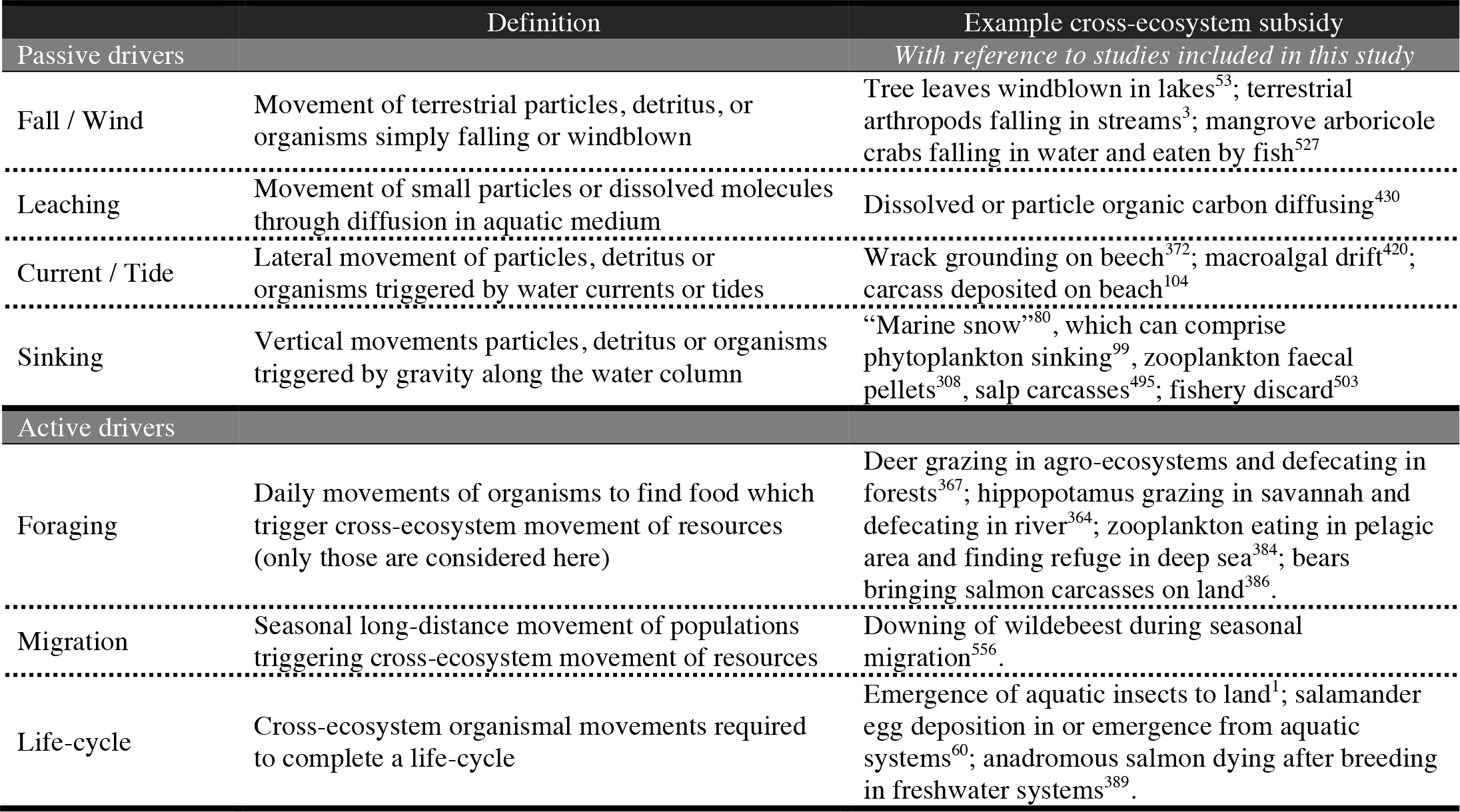
Definition of flow drivers.

**Supplementary Table 1:**
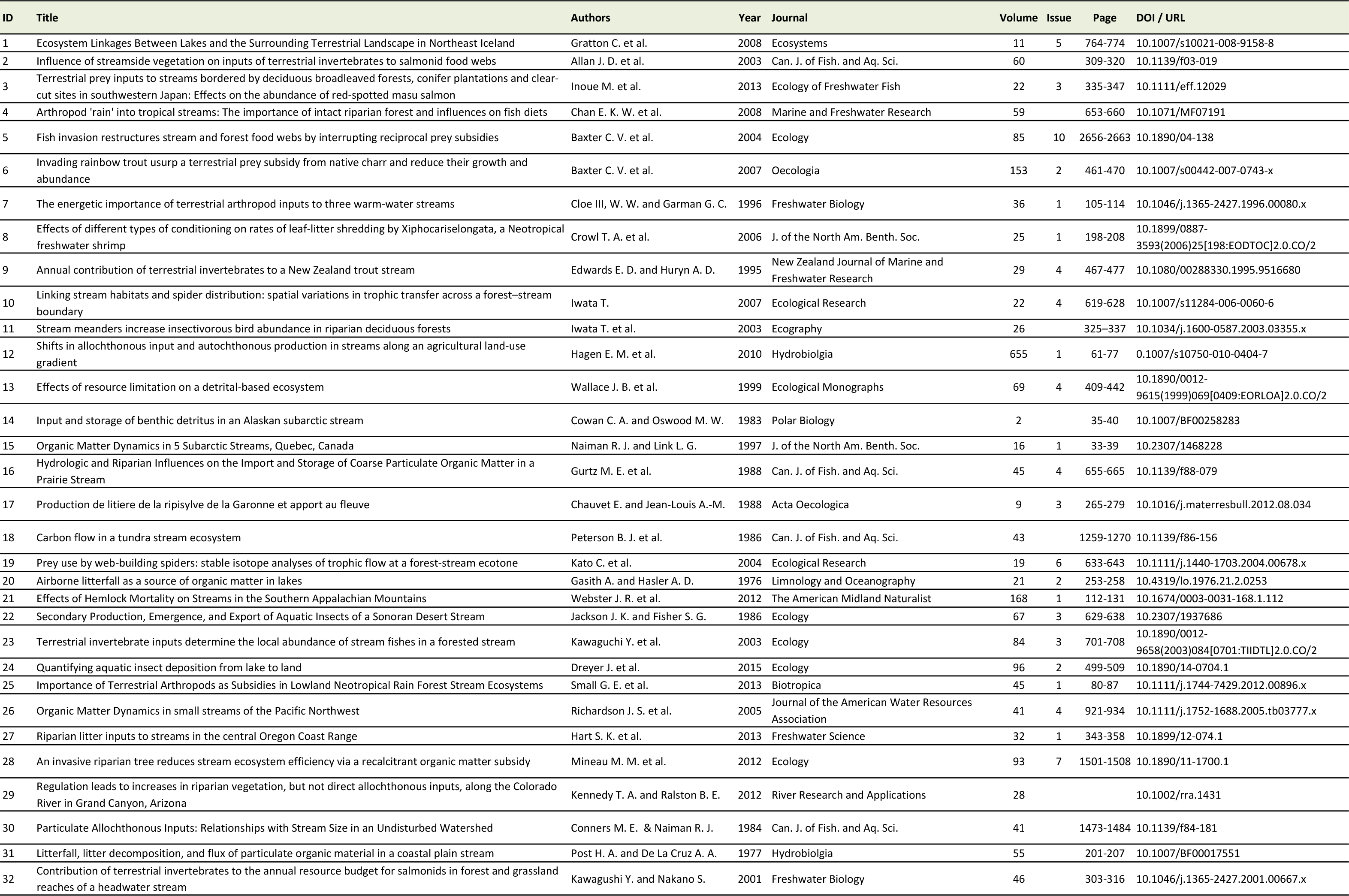
List of references from which the data are extracted; see summary of the data in Extended Data Tables 2 and 3.

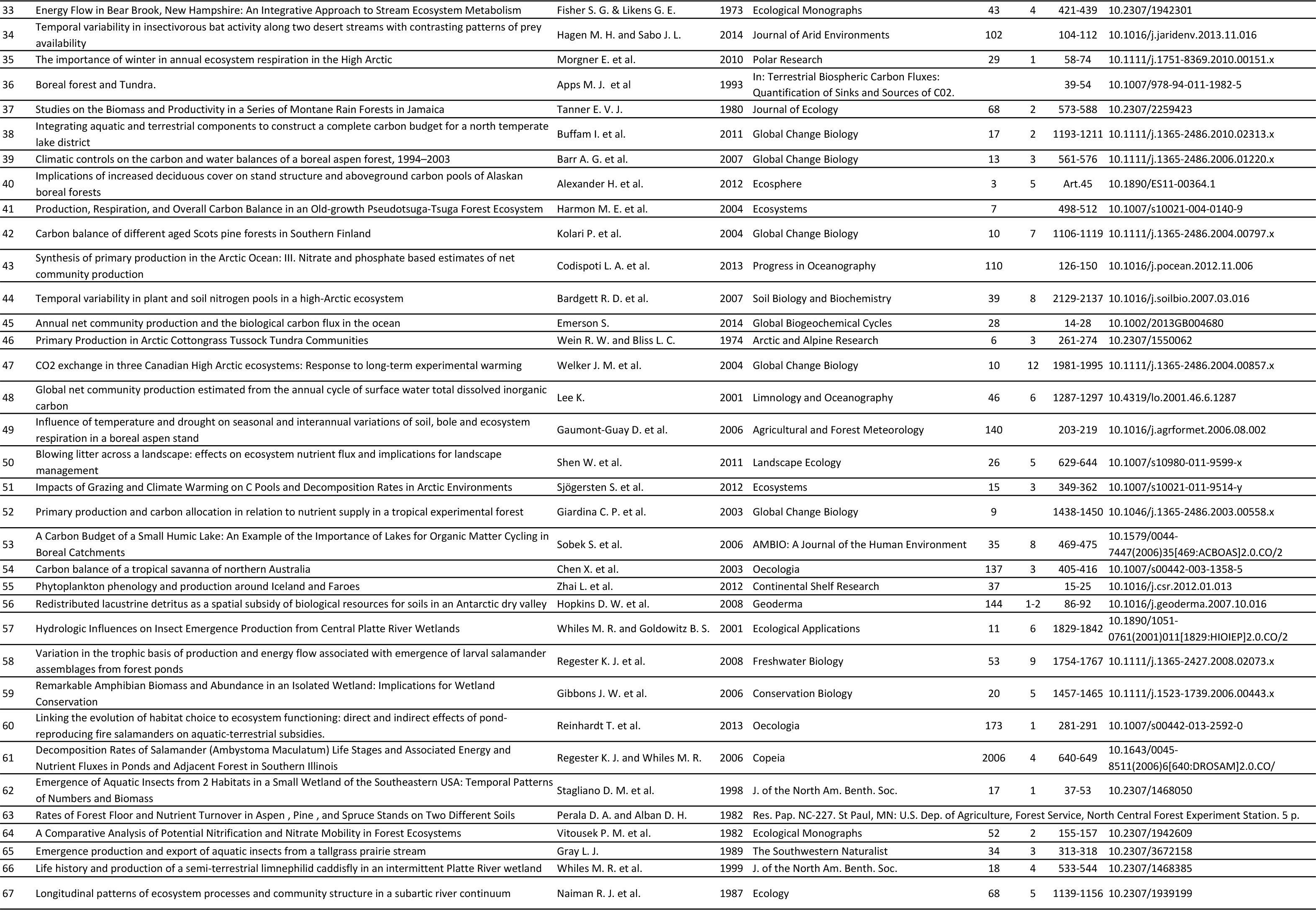

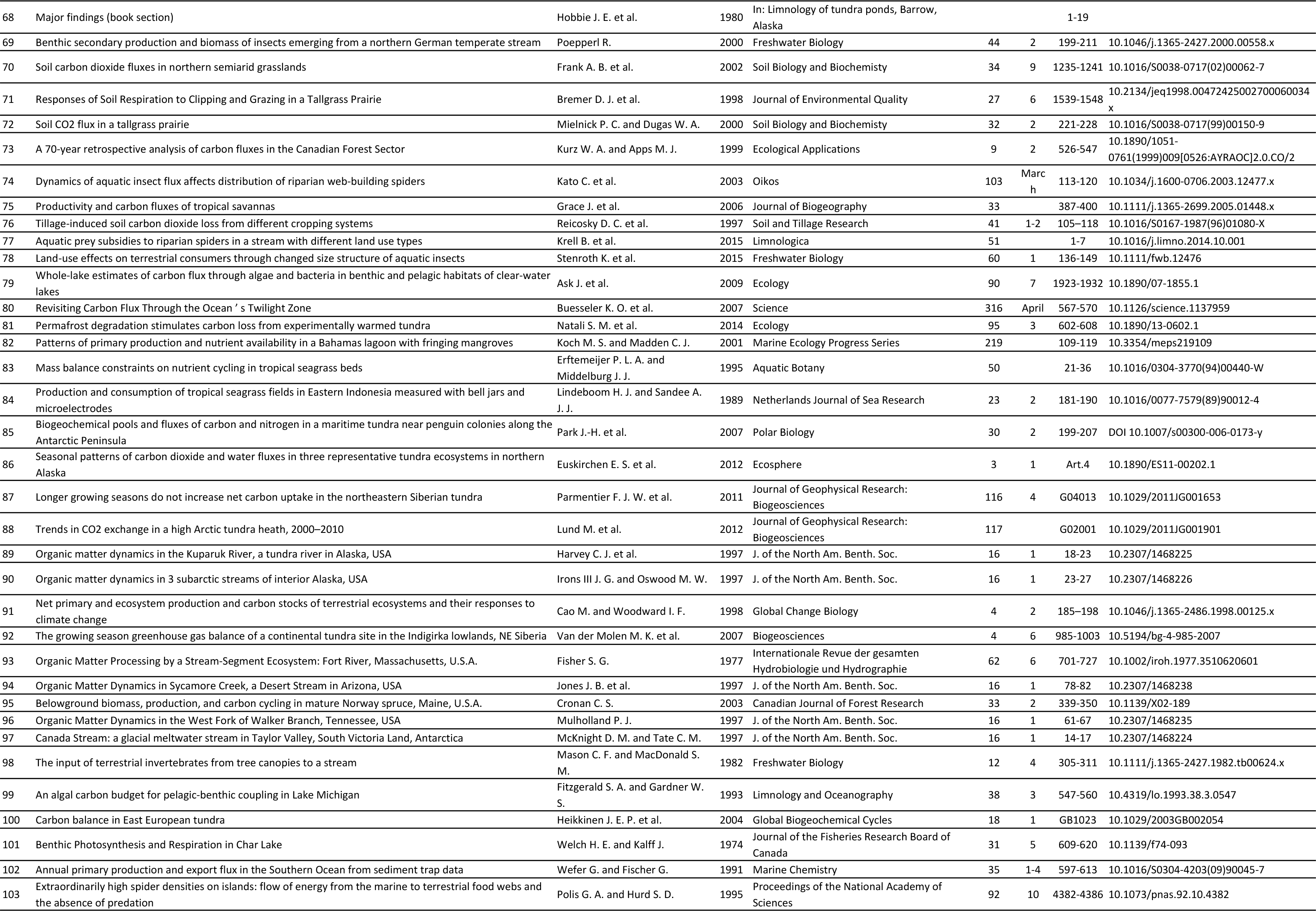

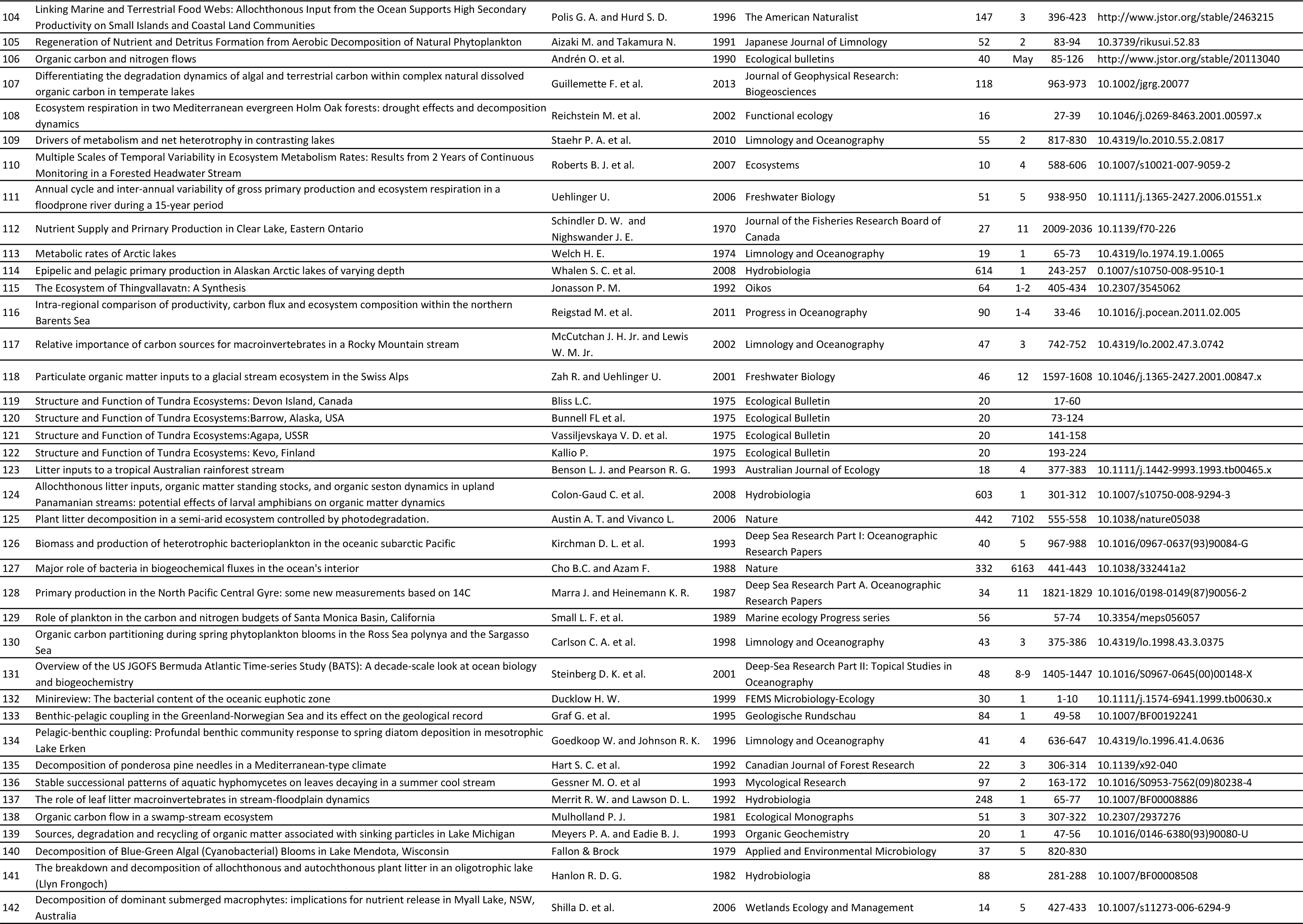

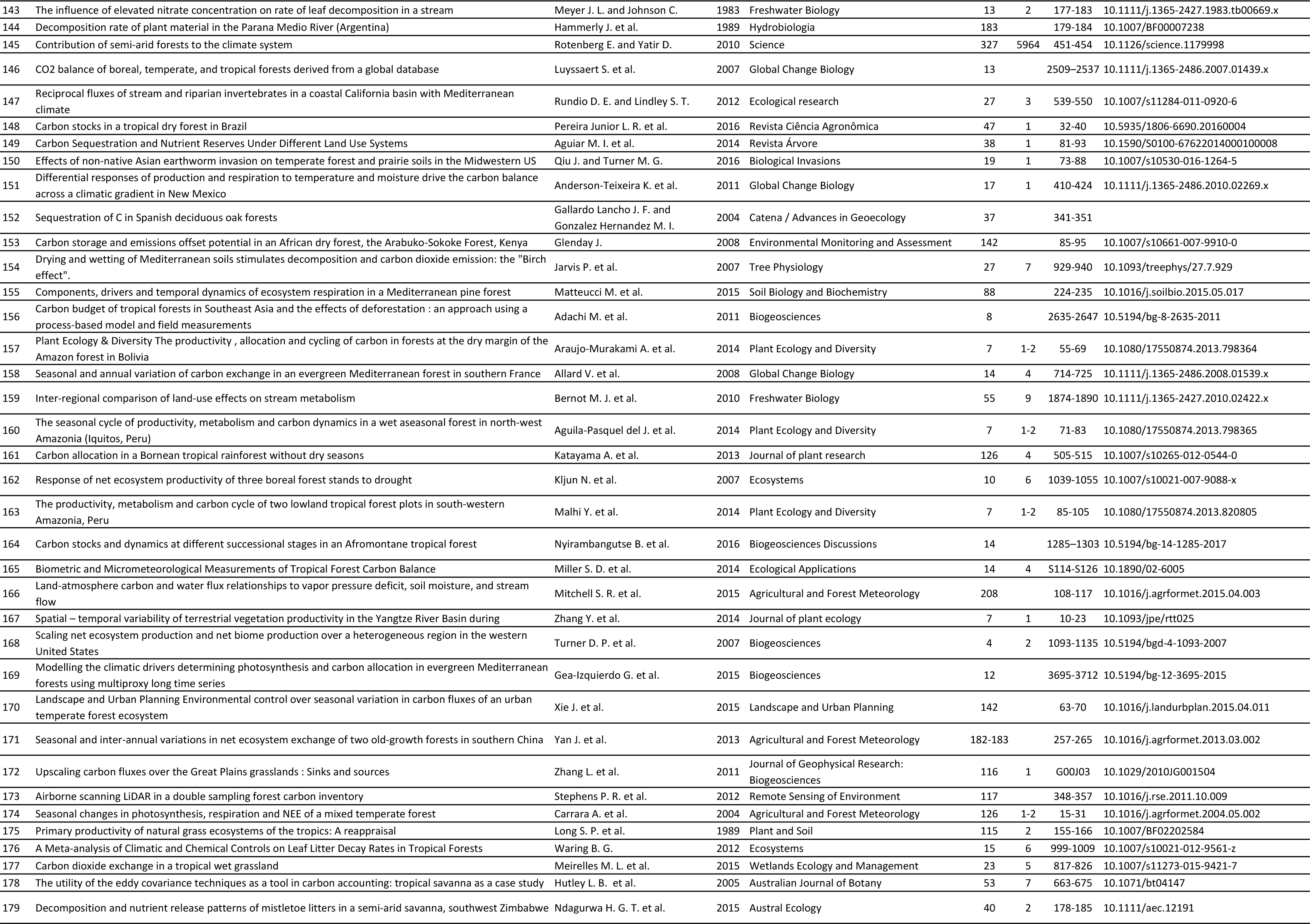

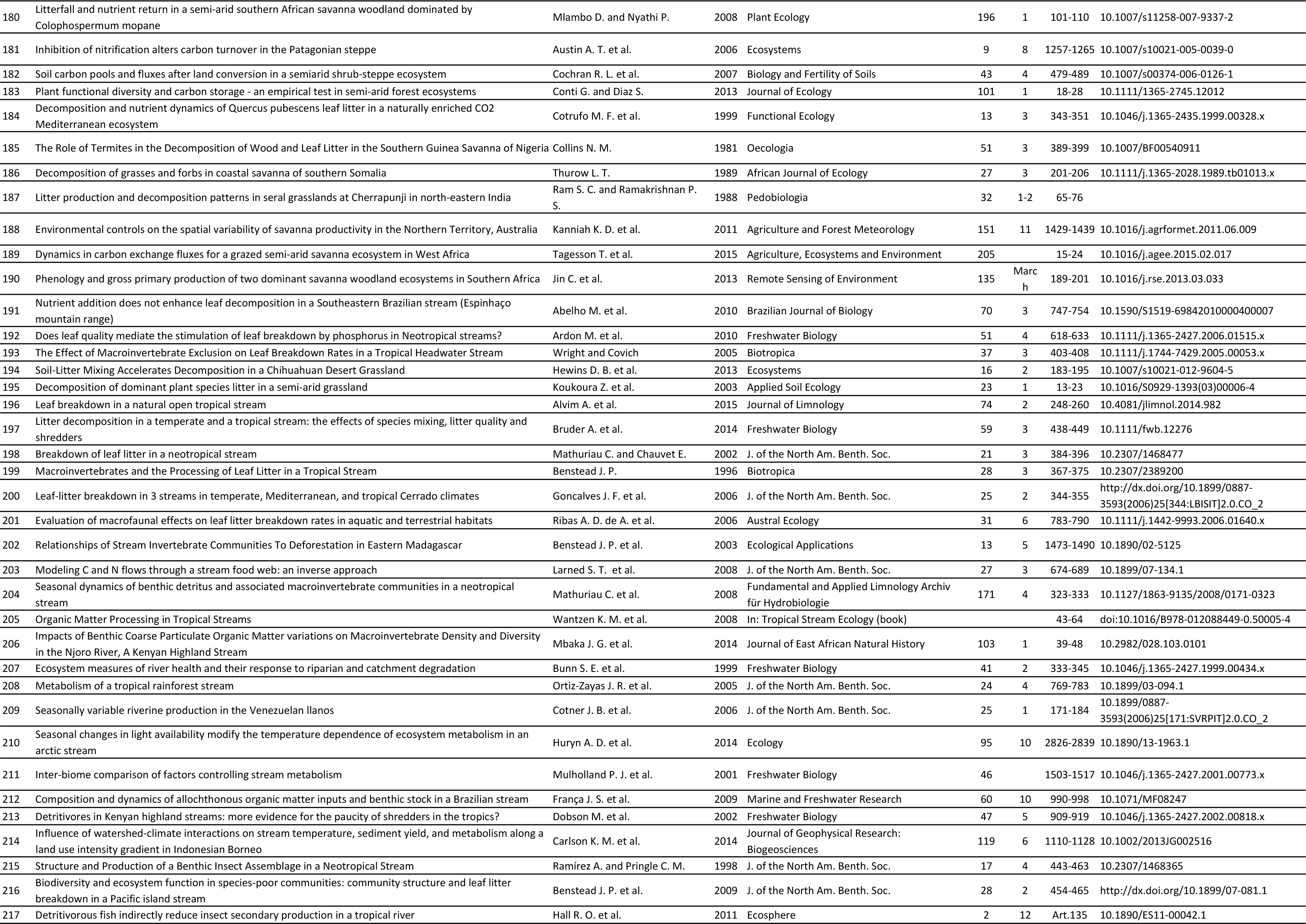

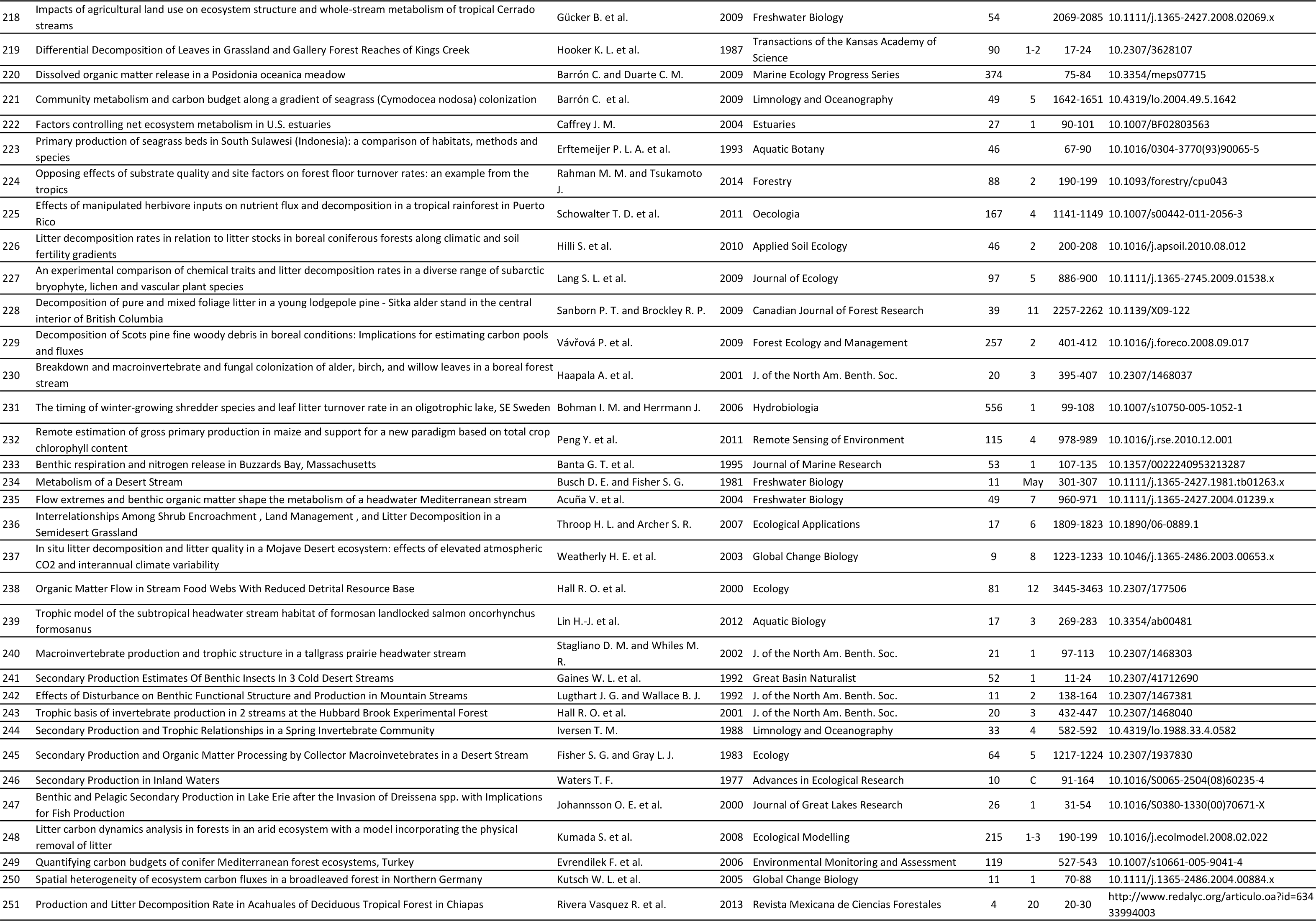

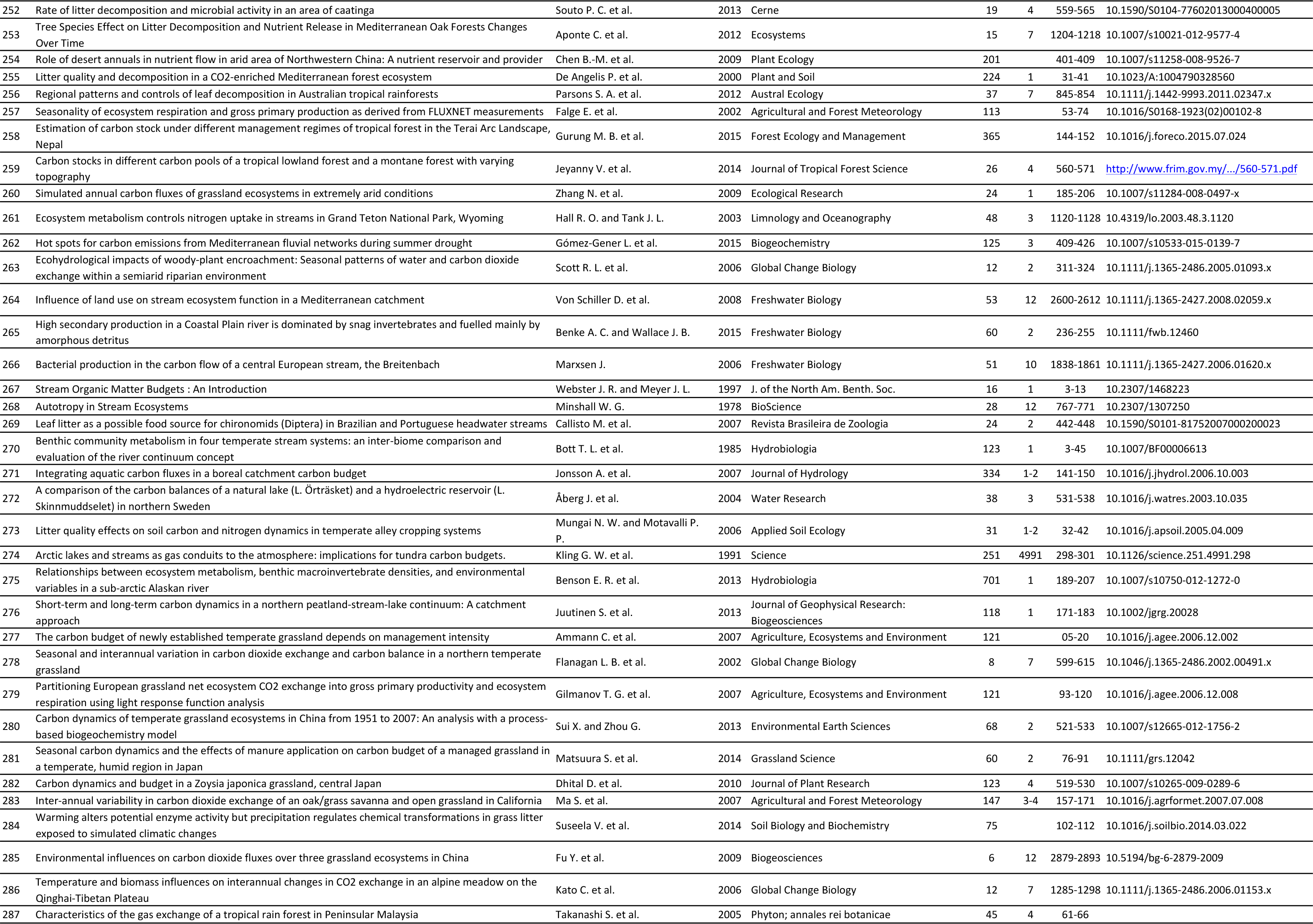

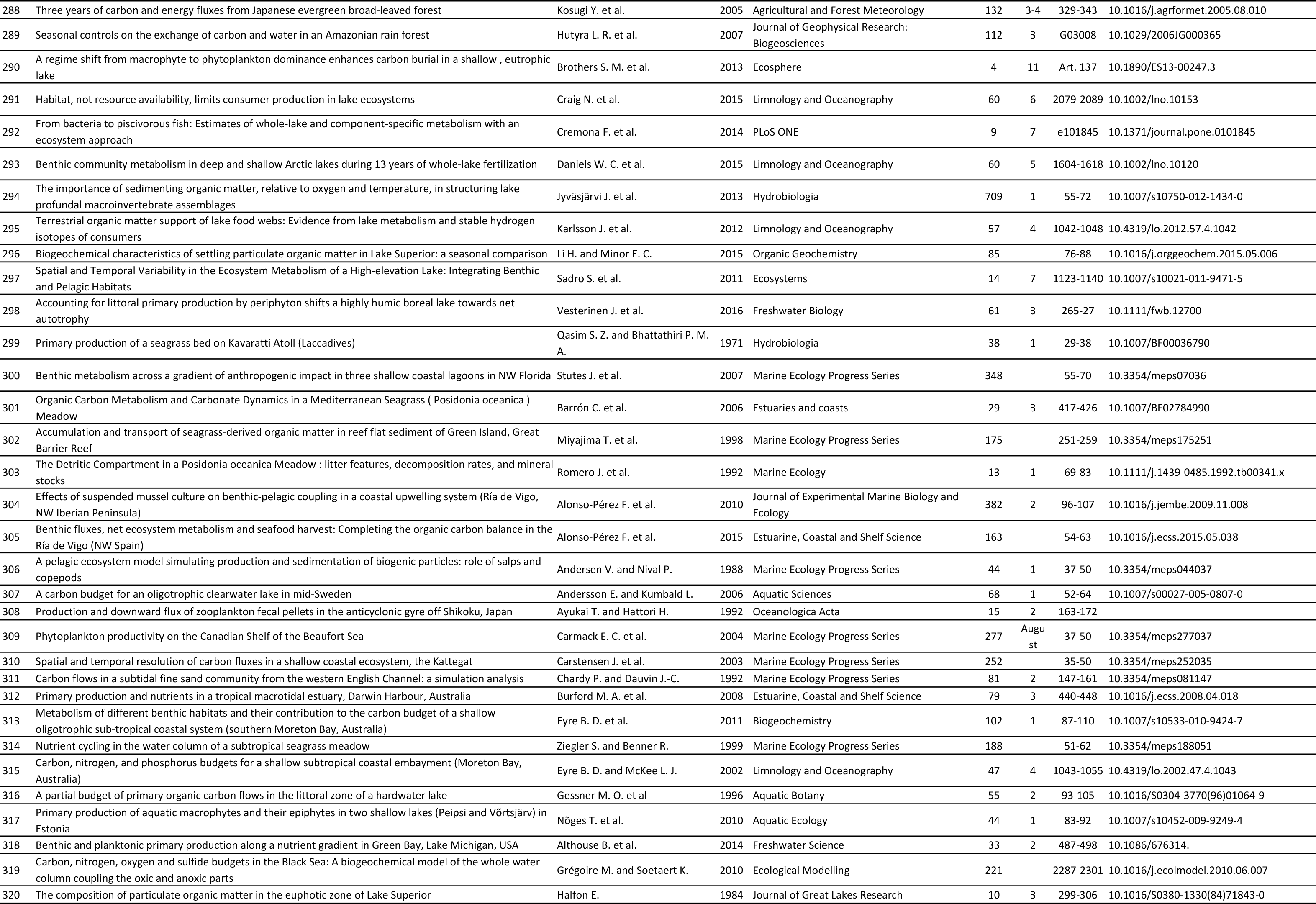

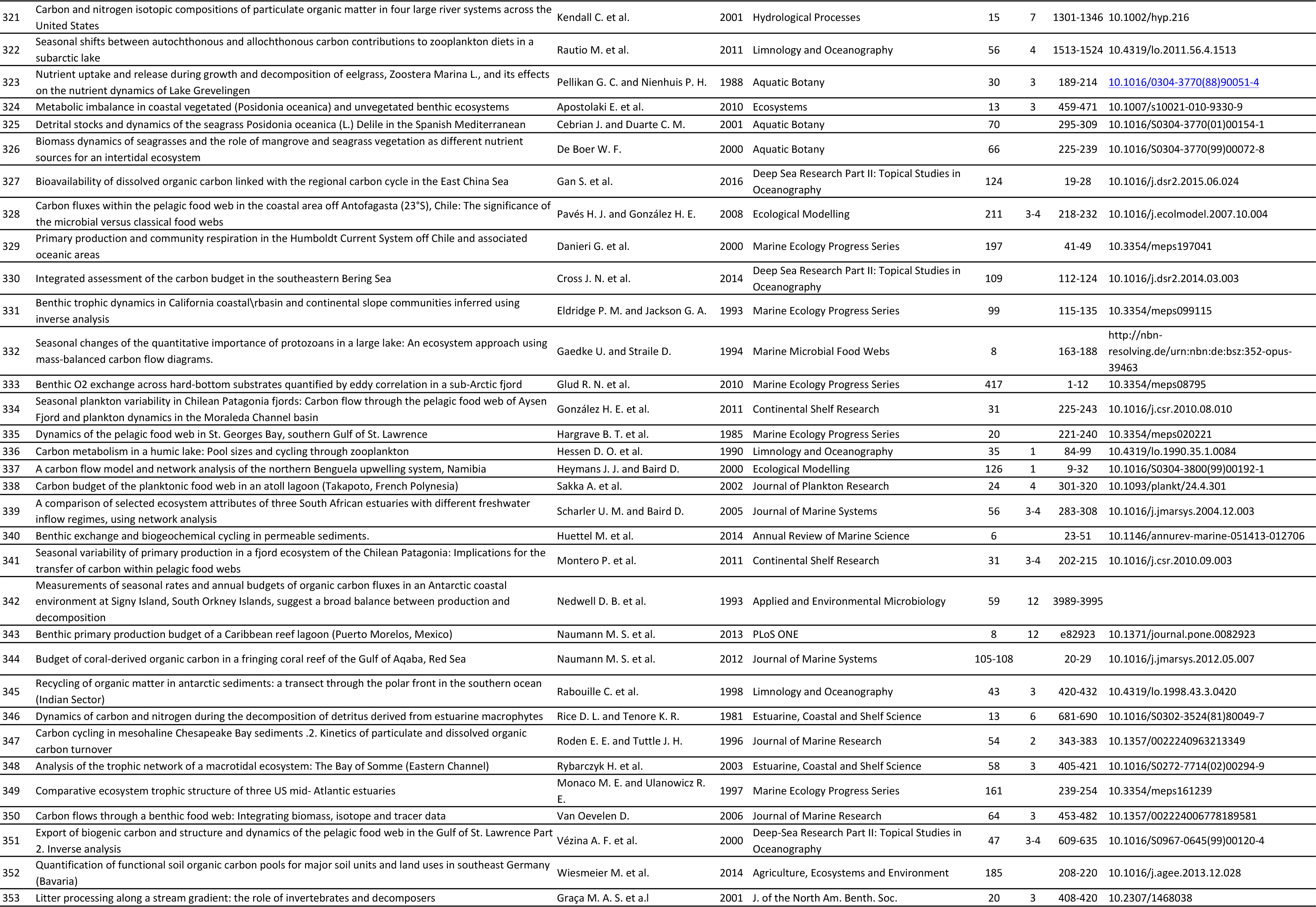

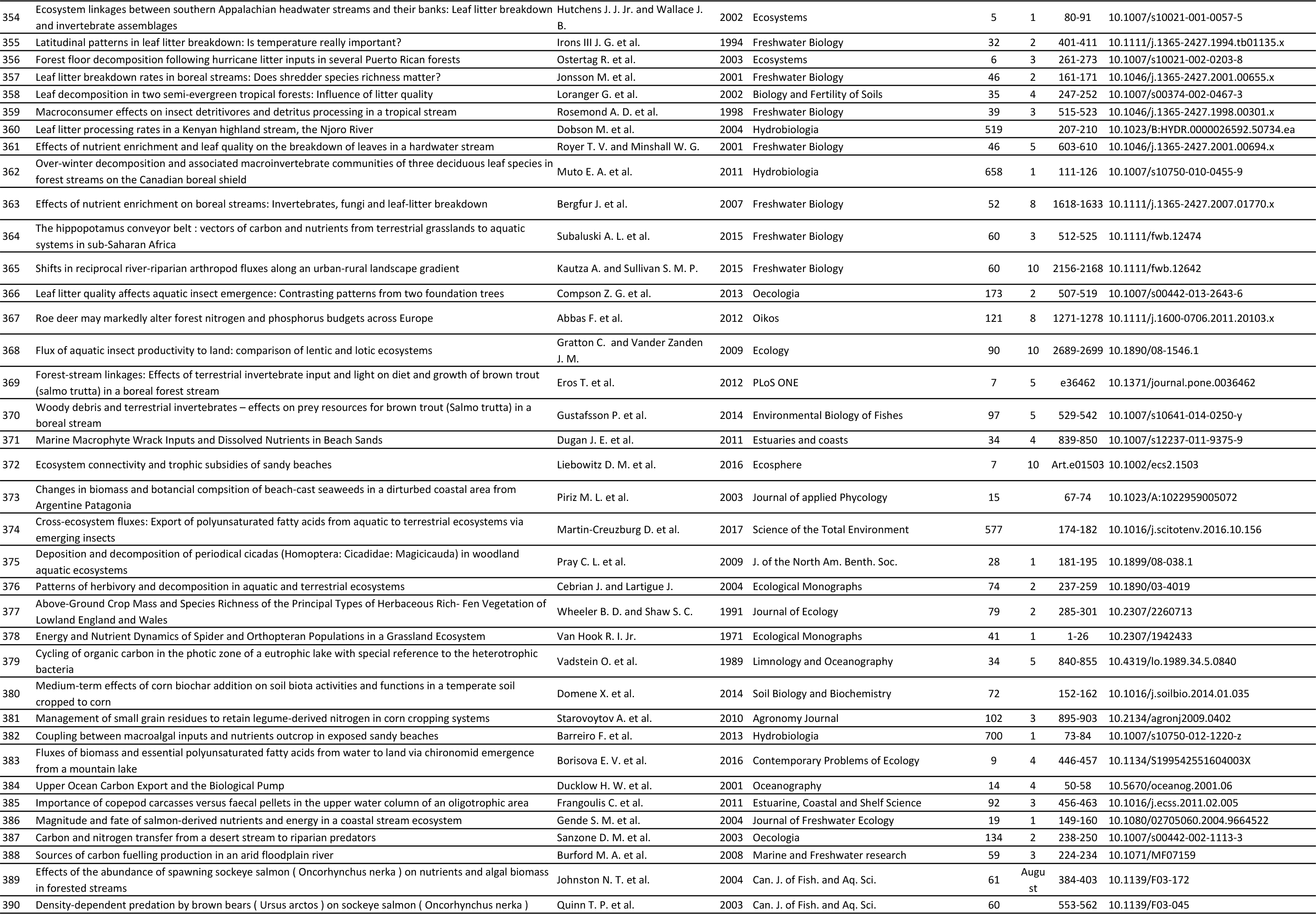

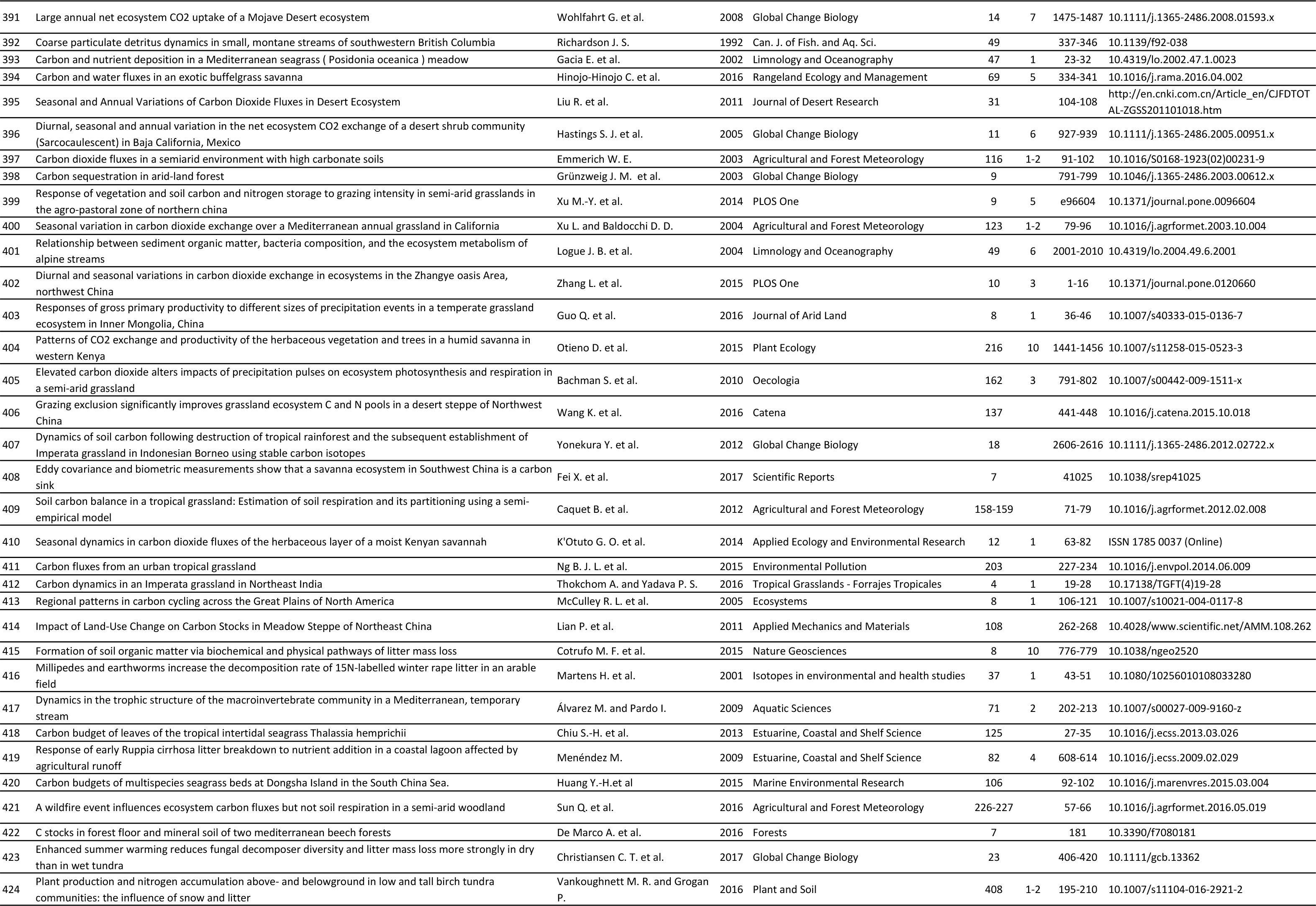

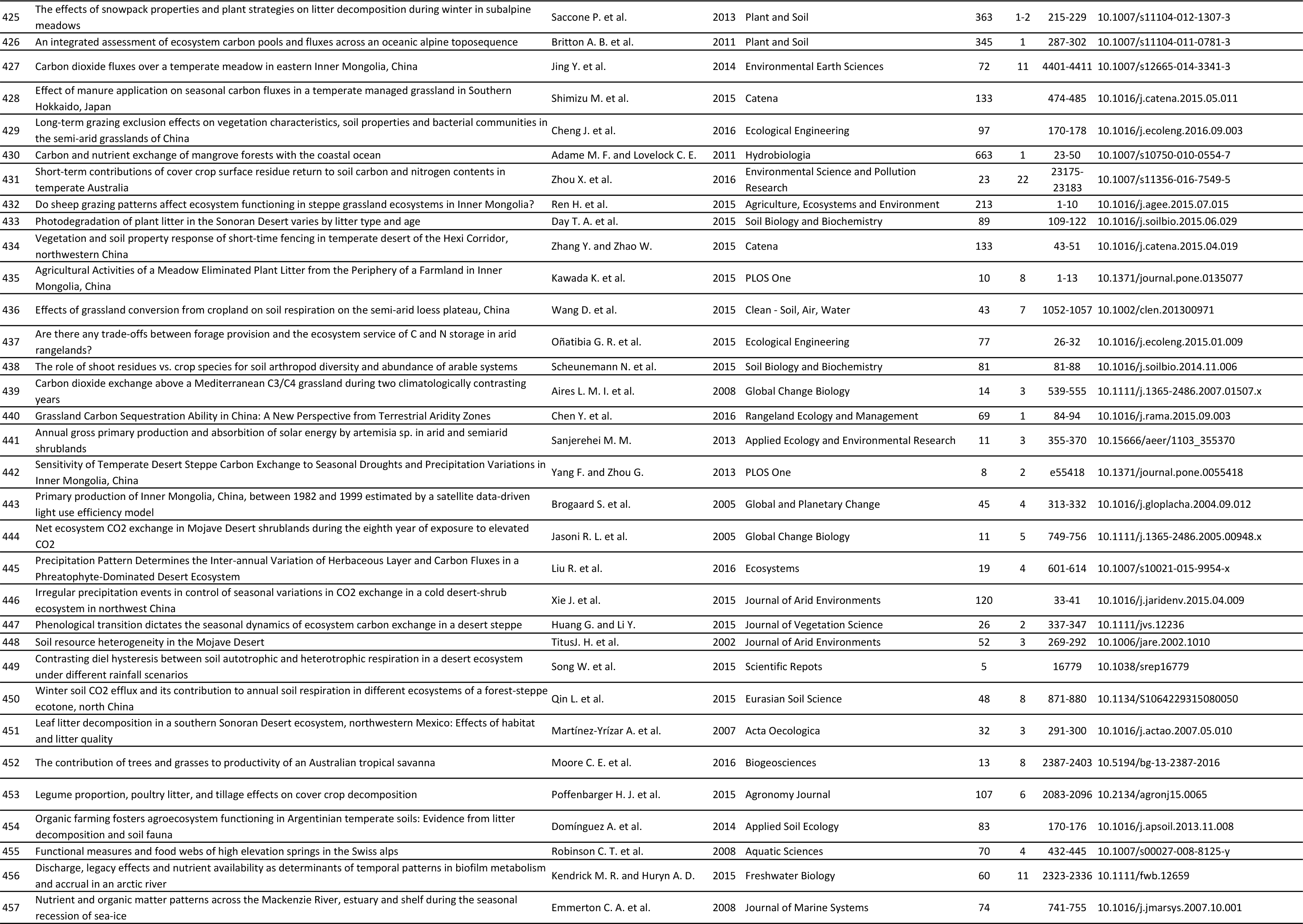

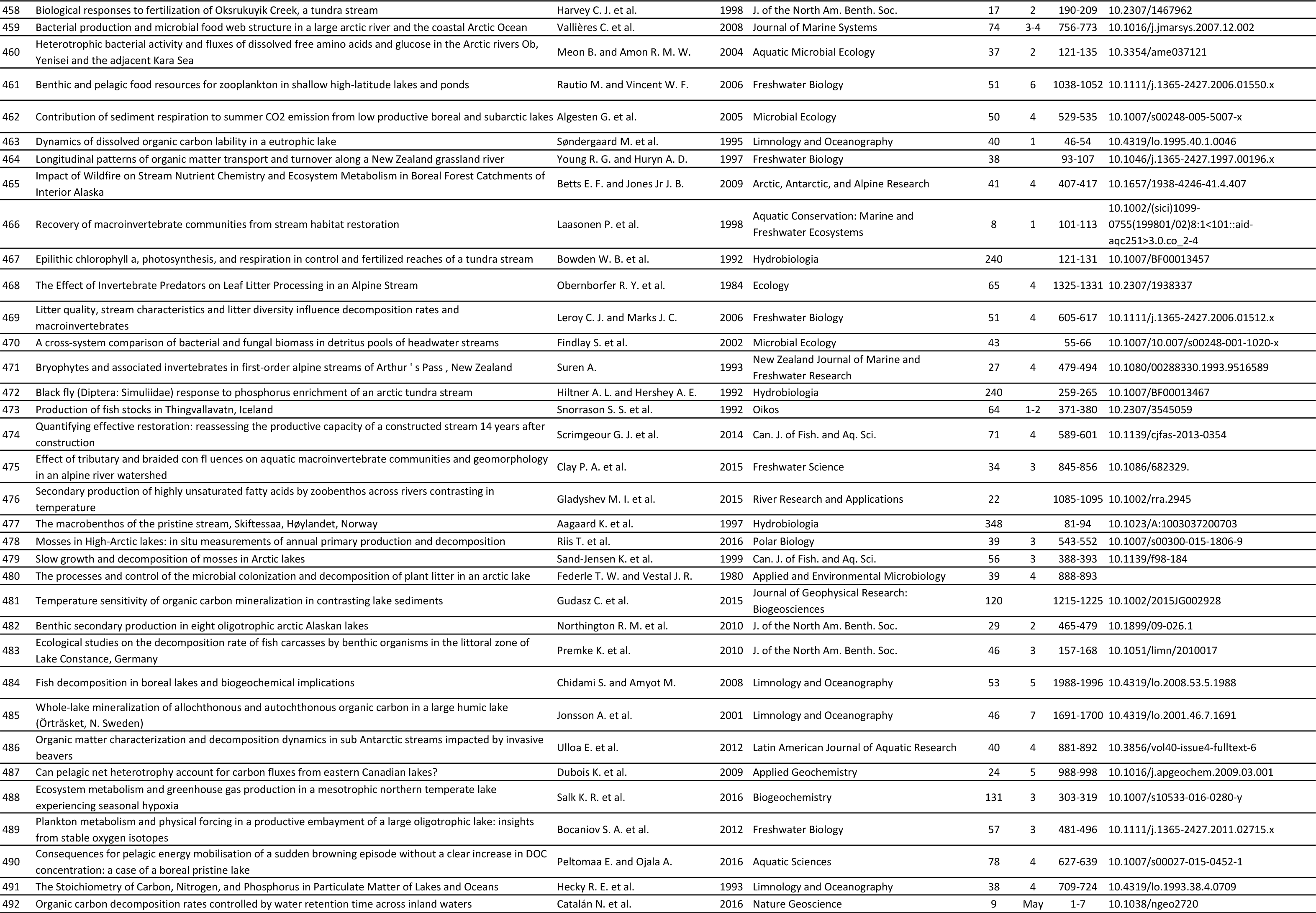

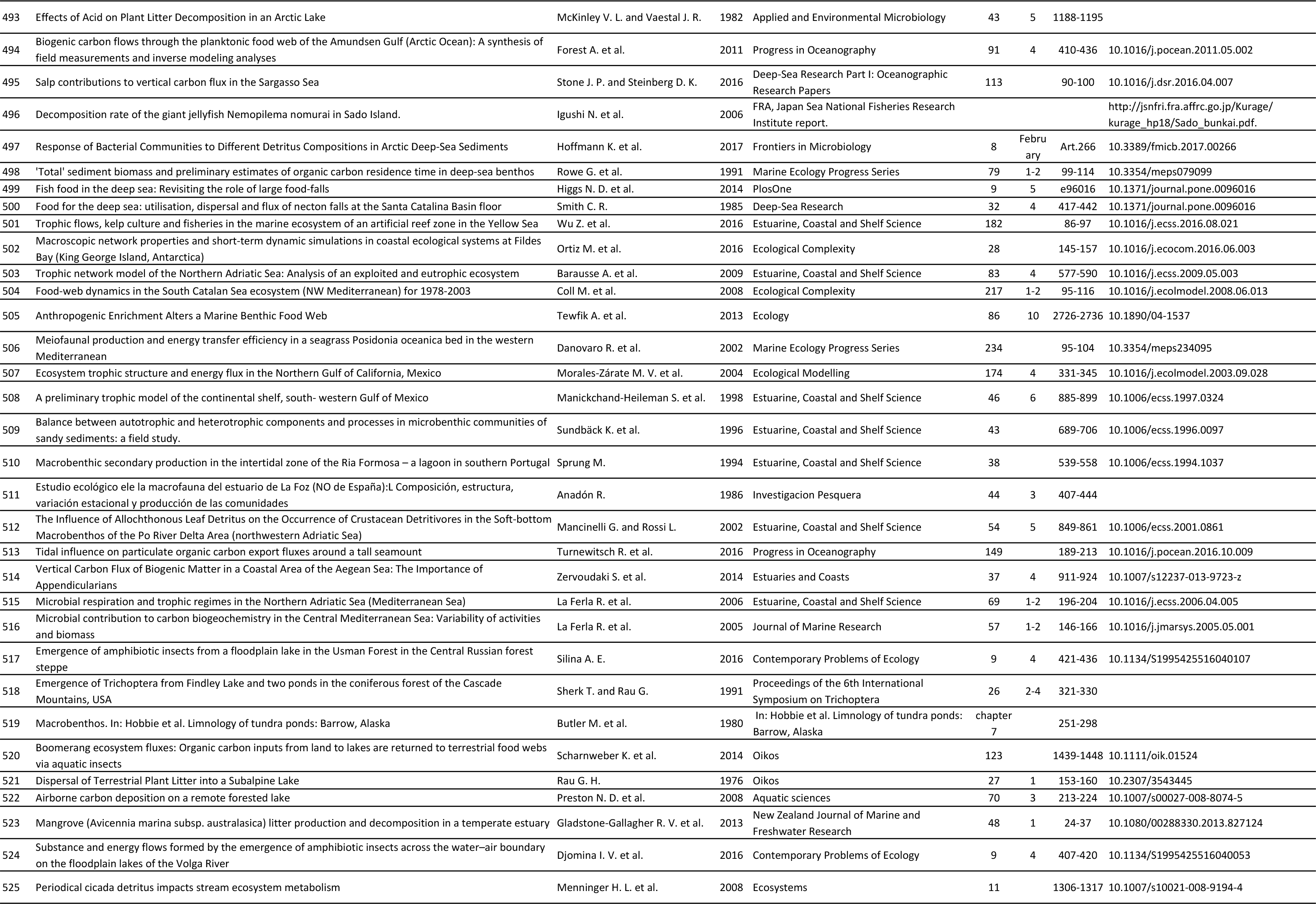

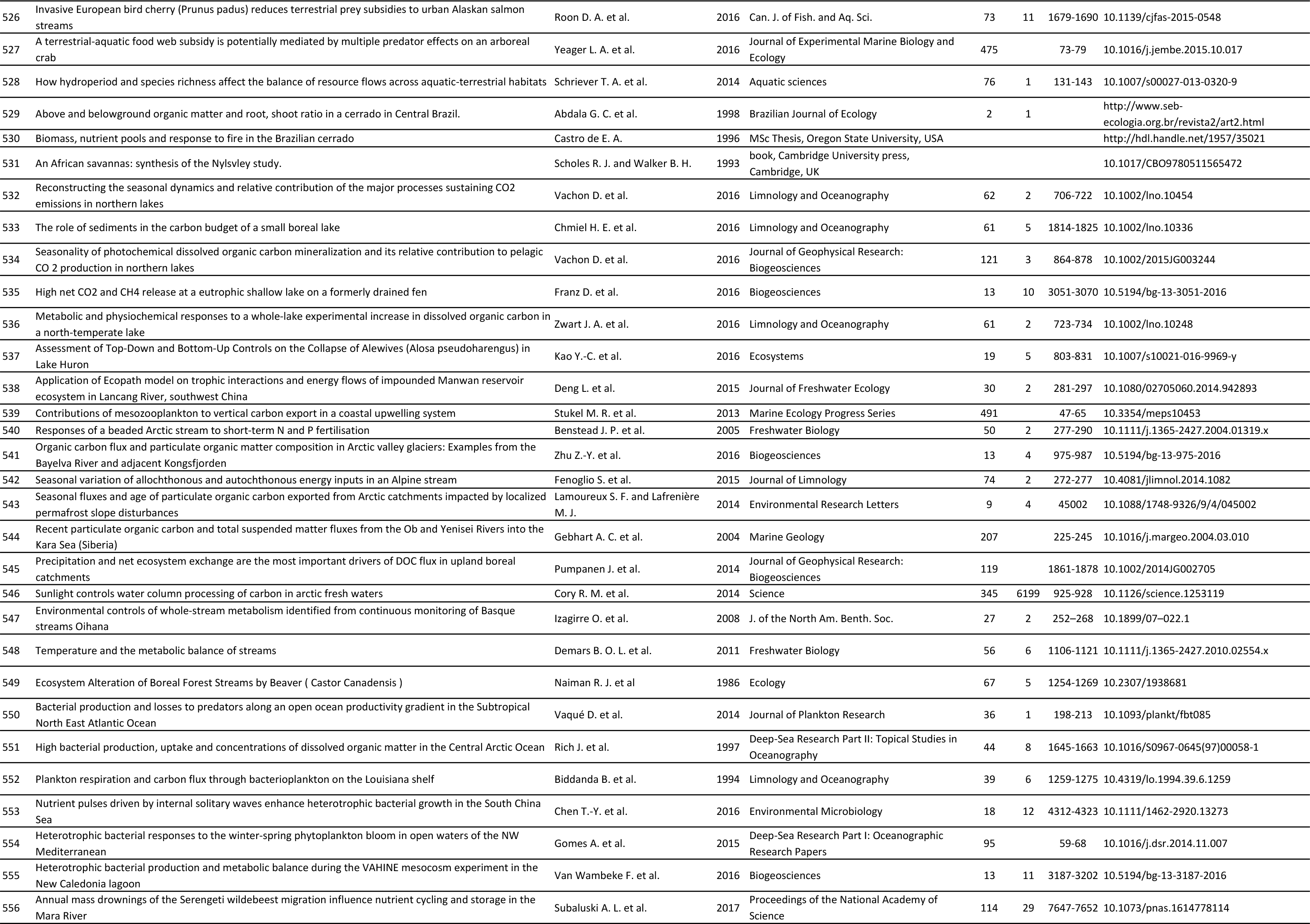

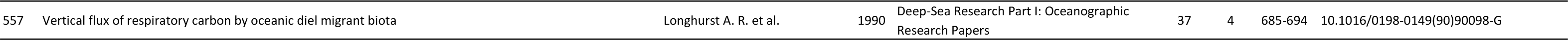

